# Freeform printing of heterotypic tumor models within cell-laden microgel matrices

**DOI:** 10.1101/2020.08.30.274654

**Authors:** Thomas G. Molley, Gagan K. Jalandhra, Stephanie R. Nemec, Aleczandria S. Tiffany, Brendan A. C. Harley, Tzong-tyng Hung, Kristopher A. Kilian

## Abstract

The tissue microenvironment is comprised of a complex assortment of multiple cell types, matrices, membranes and vessel structures. Emulating this complex and often hierarchical organization *in vitro* has proved a considerable challenge, typically involving segregation of different cell types using layer-by-layer printing or lithographically patterned microfluidic devices. Bioprinting in granular materials is a new methodology with tremendous potential for tissue fabrication. Here, we demonstrate the first example of a complex tumor microenvironment that combines direct writing of tumor aggregates, freeform vasculature channels, and a tunable macroporous matrix as a model to studying metastatic signaling. Our photocrosslinkable microgel suspensions yield local stiffness gradients between particles and the intervening space, while enabling the integration of virtually any cell type. Using computational fluid dynamics, we show that removal of a sacrificial Pluronic ink defines vessel-mimetic channel architectures for endothelial cell linings. Pairing this vasculature with 3D printing of melanoma aggregates, we find that tumor cells within proximity migrated into the prototype vasculature. Together, the integration of perfusable channels with multiple spatially defined cell types provides new avenues for modelling development and disease, with scope for fundamental research and drug development.

## Introduction

Tumor progression and dissemination are influenced through local microenvironment mechanics and degradability^1^, surface topology^2,3^, and paracrine and autocrine signaling between tumor cells and surrounding stroma^4–6^. Within this complex microenvironment, blood and lymphatic vessels play critical roles in feeding the primary tumor, while also providing an avenue for dissemination through intravasation and extravasation^7^. While simple co-culture models from transwell plates^8^, monolayers^9^, 3-dimensional (3D) spheroid co-cultures^10^, and cell-embedded hydrogel matrices^11^ have yielded great insights into tumor-stroma and vasculature interactions, considerable work remains to realize full spatiotemporal control in 3D—an essential task for understanding the functional relationships of cells, stroma, and molecular interactions in this multivariate space. And given the complexity of the signaling underlying tumor progression, creation of robust models that assemble multiple cell types in vitro has remained a challenge^12^.

In the past decade, 3D bioprinting has emerged in the tissue engineering and disease modeling spaces as a tool to attenuate the spatiotemporal properties of cells and their surrounding matrices. Recently, researchers have made advances in spatial organization of in vitro tissues by printing into support baths of suspended microgels^13–18^. These support baths fluidize under shear force as the microgel particles near the print nozzle translate around the tip, while subsequently supporting the ink that is deposited. This enables the freeform printing of inks in all dimensions, allowing for complex structures and taller prints^19^. Concurrently, Lewis and colleagues created the first method for directly writing vasculature through sacrificial inks to create vascularized hydrogels^20,21^ and they have further extended this work to printing into a support bath of organoids to form thick vascularized tissues^22,23^. Yet, translating direct writing of vasculature into microgel support baths to create endothelial lined channels in the presence of live cells has not previously been reported.

Here, we present freeform vascular printing in cell-laden microgel suspensions where a sacrificial ink deposited within photocrosslinkable microgels defines hollow channels amidst printed cancer and stromal structures. The platforms advance lies in its modularity, in that virtually any combination of cells and cell structures to be spatially defined within the suspension with controlled proximity to channels. As proof of principle, we spatially organized three important contributors to tumor progression: (1) fibroblasts dispersed uniformly within the microgel suspension, (2) primary tumor cells and structures in defined 3D architectures, and (3) endothelial cells within interpenetrating hollow channels towards perfusable vessels (**Figure 1A**). Further microenvironment control is afforded by changing the microgel composition and chemistry, facilitating tunable local and global mechanics of the microgel construct.

**Figure 1.**
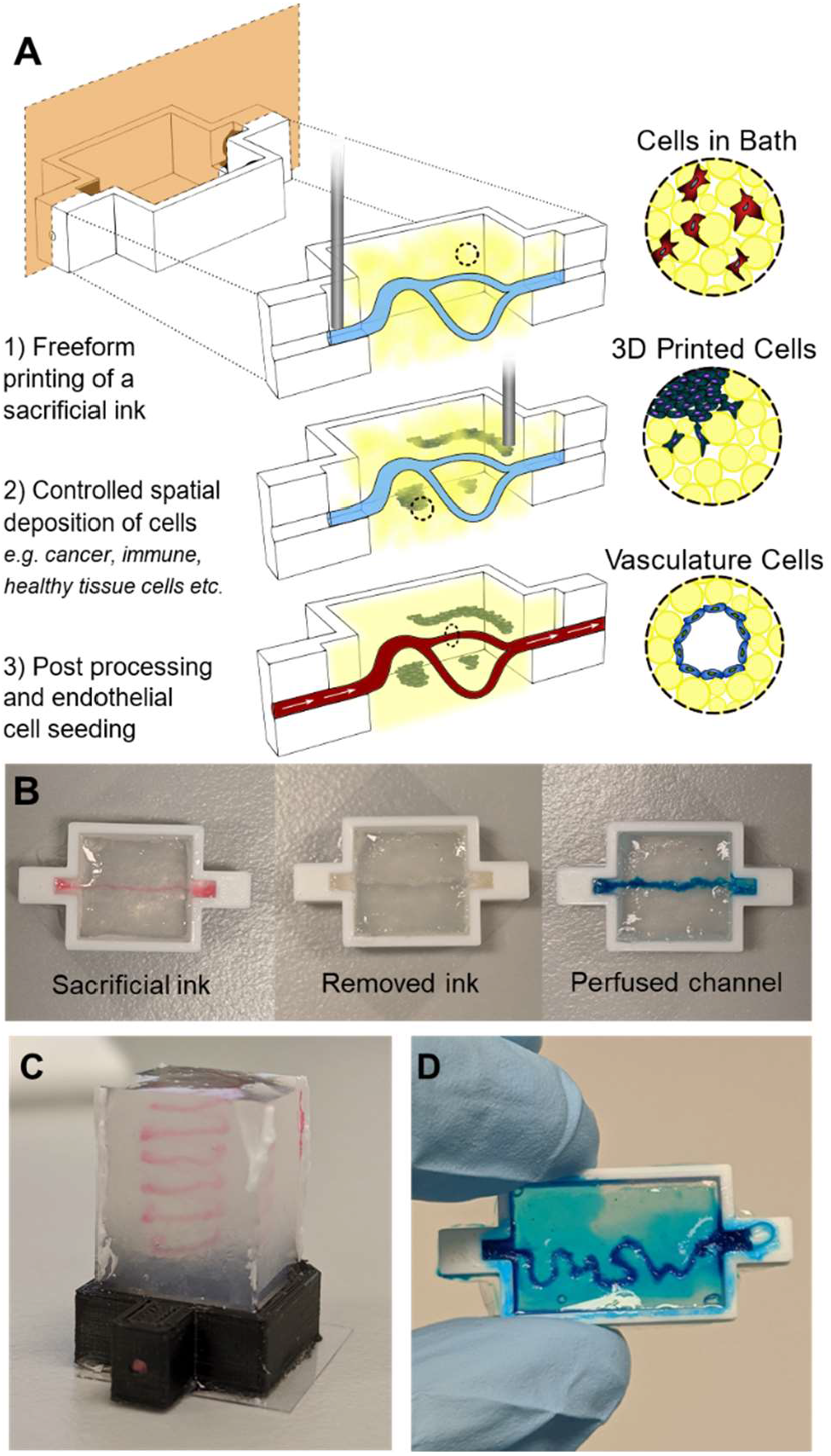
Photocrosslinking to stabilize suspension microgels allows for complex organization of cells in three distinct ways. a) (1) An uncrosslinked microgel suspension, with or without cells, is placed in a reactor where a sacrificial ink is freeformly printed. (2) More cell types can further be printed as different shapes and sizes at various proximities to the sacrificial ink. (3) The suspension is photocrosslinked followed by removal of the sacrificial ink and subsequent seeding of endothelial cells on the hollow channel walls. b) Macro images of the three stages of hollow channel formation: printing of the ink, photocrosslink and evacuation of the ink, and perfusion of the hollow channel for seeding. c) A macro image of a 7mm tall spiral print of Pluronic F127 ink in microgel suspension. d) An image of blue dye that has been perfused through the letters “UNSW” that were printed and evacuated.

## Results

In contrast to dissolvable gelatin microparticles used in previous work^15,16^, we synthesized gelatin-methacryloyl (GelMa) microparticles using a water-in-oil emulsion; liquid GelMa is added dropwise to 40°C oil under stirring followed by cooling to 10°C to physically crosslink the microparticles, leaving methacryloyl moieties for further crosslinking. Adding acetone then dehydrates the microparticles and allows for easy washing and weighing. When rehydrated, the microparticles have an approximate diameter of 100 microns (**Figure 2A**). Since yield stress fluid properties can vary greatly with small changes in suspension compositions, we weighed and hydrated our dried microparticles with consistent particle to liquid ratios. These suspensions were rested for at least 24 hours prior to use since acetone dried GelMa can take days to rehydrate (Figure S1). At hydrated volume fractions of ∼50- 60+%, the microparticles reach a jammed state where they lock in place by frictional and repulsion forces^19^. These jammed particle suspensions behave as a solid under equilibrium conditions but will flow like a liquid once a critical shear force is applied. Swelling tests of the GelMa microparticles showed they rehydrate to ∼10x their dried weight, which we used to hydrate our suspensions to the target 60% volume fraction of particles consistently.

**Figure 2.**
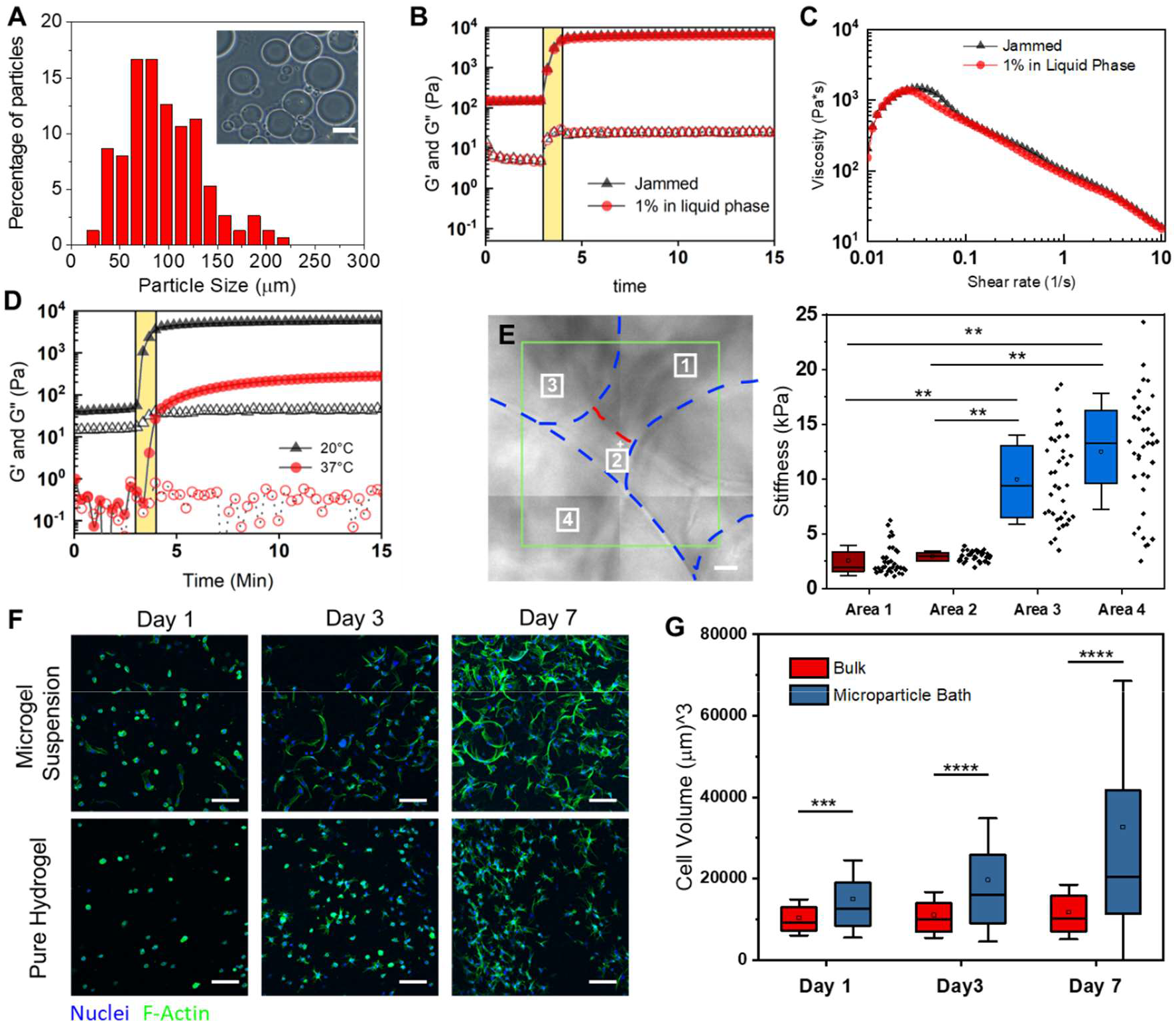
Characterization of GelMa microparticle support suspensions for 3D printing. a) Size distribution (n=100) of the 10 wt% GelMa microparticles. Inset image is a representative optical image of the microparticles. b) Rheological analysis of the gelation of suspensions with (red markers) and without (black markers) a 1 wt% GelMa filler in the liquid phase. Closed markers are the storage modulus (G’) and open markers are the loss modulus (G”). c) Uncrosslinked microgel viscosity as a function of Shear rate for suspensions with (red markers) and without (black markers) filler. d) Effect of melting a microgel suspension (red markers) on the storage modulus (G’) when compared to unmelted heterogenous suspension (black markers). e) Contact mode atomic force microscopy (AFM) over 4 regions of a suspension bath surface. Force curves (n = 36 per square) were taken at each 10μm x10μm region of the surface where the analyzed young’s modulus is plotted for each region’s curves. f) Z-stack Projections (100μm z-stacks of 50 slices) of ADSCs stained with Hoechst(blue) and Phalloidin (green) in microgel suspensions and a pure hydrogel (10 wt% GelMa) g). Scale bars: 10μm (e), 100μm (f).

By functionalizing the gelatin with methacryloyl groups, our microparticle suspensions can be chemically crosslinked within and between the particles to stabilize the matrix. After exposure to 60 seconds of 395nm light on a rheometer, the microgels gain a 2 order of magnitude increase in storage modulus (**Figure 2B**) and become stable under shear forces (Figure S2A). To gain greater control of the local mechanics of the gel as well as aid in printability, we added a fraction of soluble GelMa to provide a means of effectively “stitching” the jammed suspension together after printing. By hydrating the dried particles with a liquid solution of low weight percent GelMa, the same bulk rheological properties of the microgel suspension can be maintained, while now creating a soft matrix around the stiff particles. When hydrating the particles to 40% volume fraction with a 1 wt% GelMa solution as the liquid filler, we achieve near identical bulk mechanical properties to the jammed microparticles (no filler) while changing the interstitial space from pure liquid to a soft matrix (**Figure 2B**). When applying a shear strain rate sweep on the suspensions, both exhibit similar yield stress fluid properties—each demonstrating high printability. Suspension with fillers also demonstrate stability under shear forces once photocrosslinked (Figure S2A). Decreasing the light exposure had minimal effect on the gel strength of filler suspensions with a <1% decrease in storage modulus; however, increasing light exposure to 120 seconds gave a 23% increase (Figure S2b). Stabilized suspensions warmed to 37°C where the physical crosslinks release had a drop in strength of only 7% (Figure S2C), demonstrating stability of the network through the covalent modifications. The bulk mechanical properties can also be tuned by varying the weight percentage of the GelMa used to make the microgel suspension, where particles formed with 15 wt% GelMa formed suspensions with a 70% higher storage modulus (Figure S3).

It is well appreciated that both global and local mechanical environments can play a large role in directing cell function and behavior^24,25^. Therefore, multiple classes of mechanical testing are required for this type of heterogeneous microgel suspension. To highlight this heterogeneity, we melted a suspension solution prior to photocrosslinking and found a decrease in strength of nearly two orders of magnitude (**Figure 2E**). We further demonstrate this heterogeneity with AFM force curves (1μm radius spherical borosilicate probe, 36 curves per 10μm x10μm regions) taken at 4 different locations across the surface of our crosslinked microgel. The regions over the microparticles have relative moduli of over 5 times that of the filler regions. However, even over the stiffer particles, there is wide variation at the local scale as the filler material wraps itself not only between particles, but around them as well, creating broad variability in stiffness the cell experiences. This contrasts with similar microporous particle scaffold (MAP) systems that contain discrete pockets of heterogeneity^19^.

We next set out to explore the exciting possibility that our printing support matrix would be beneficial to integrated live cells. We began by seeding adipose derived stem cells (ADSCs) at one million cells/ml of microgel suspension. Initial live/dead staining of cells indicated high cell viability (Figure S4). However, the nature of the scaffolds made it difficult to image samples thicker than 0.5mm. The particles have a much higher index of refraction compared to the filler phase, thus leading to significant light scattering during imaging. To circumvent this caveat, we adapted our recently reported optical clearing technique^26,27^ where index matching allows increased imaging depth with minimal light scattering (Figure S5). To evaluate our clearing and imaging protocol, we loaded one million ADSCs per ml into our suspension, with a pure bulk GelMa matrix of comparable mechanics as a comparison. Given the porous nature of the microgels, and the tendency for cells to spread anisotropically in 3D, we sought to compare cell volume and surface area in 3D rather than with 2D projections. High resolution z-stacks of cells stained with phalloidin and Dapi were imported into Imaris to segment the cell and nuclear volumes (Figure S6A). There are increasingly significant differences of cell volume between our suspension and bulk GelMa on days 1, 3 and 7 (Day 1 p <0.001; Day 3,7 p = 0.0001) with the suspension cells increasing volume by ∼100% from day 1 to 7 (p = 0.001) (**Figure 2G-H**). While no significance difference in cell volume was found across any days for the bulk gels, cell surface area measurements show a ∼25% increase from day 1 to 3 (p = 0.024) and a ∼50% increase from day 1 to 7 (p = 0.001) (Figure S6B). Over time, the cells within the microgels took on the microporous architecture and proliferated, suggesting both viability and bioactivity of the interconnected network.

To vascularize our gels, we designed and 3D printed plastic molds with inlets for aligning needles for removal of the sacrificial ink and the seeding of vascular cells (Figure S7a). Since our scaffolds melt at physiological temperatures, we needed a sacrificial ink that liquifies as its temperature is lowered. Pluronic F127 was chosen as Lewis and colleagues have shown great success using this material in direct writing due to the tunable lower critical temperature^21,28,29^. Initially, we found that if the Pluronic F127 was not fully solidified, it would swell with water as printed, begin diffusing apart, and not anchor as it was printed. Therefore, we used 29 wt% Pluronic F127 for defining our vasculature channels because it fully sets at our laboratory’s ambient temperature (19°C) (Figure S7B). Red and blue dyes were added to the Pluronic ink to aid with visualization. Troughs were added to our vascular printing reactors to facilitate ink removal and cell seeding (**Figure 3A**). After ink deposition, the suspension is photo crosslinked and the microgel is placed in the refrigerator(4°C) for 10-15 minutes to liquify the Pluronic F127. The ink is then removed by syringe, leaving a hollow channel inside the microgel matrix (Figure S7E).

**Figure 3.**
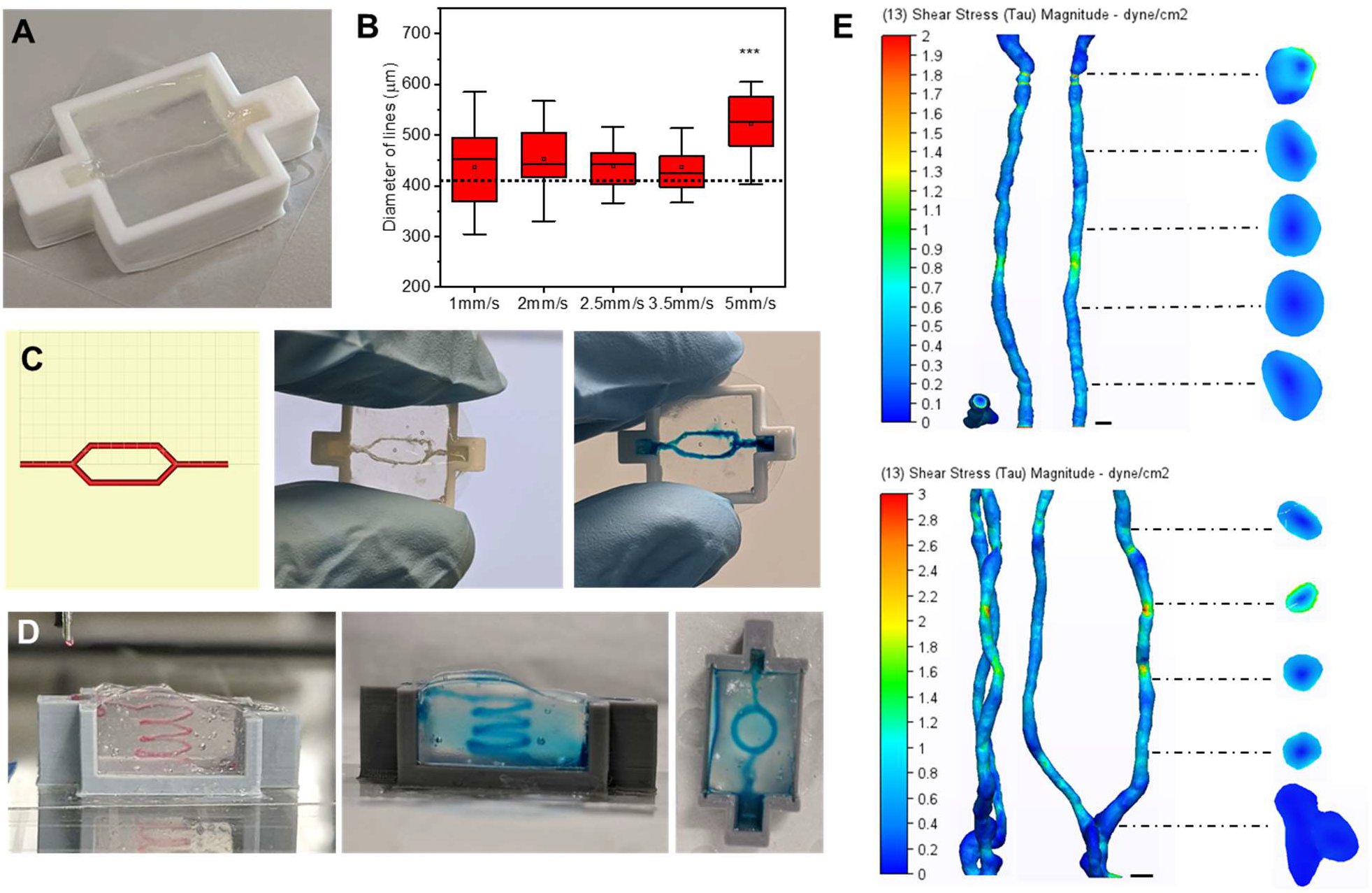
Freeform printing of complex perfusable channels. a) An image of a hollow channel in the PLA 3D printed reactors used for vascular printing. b) Box plots of diameters of the Pluronic F127 ink (n=28-35) printed into suspension baths at different speeds (5mm/sec compared to 1,2,2.5, & 3.5mm/sec; p = 0.0101). c) Images of the print head tool path for a bifurcation in Pronterface (left) followed by an evacuated bifurcation print (center) and the same print perfused with a blue dye (right). d) Images of a four-loop spiral printed with Pluronic F127 (right) that was evacuated and perfused with a blue dye (center and right). e) Wall shear stress heat maps for computation fluid dynamics (CFD) simulations run on representative single (top) and bifurcated (bottom) channels imaged via MicroCT. Scale bars: 400μm.

To create a system where consistent printing and reproducibility is ensured, we optimized printing parameters through design of experiments using print speed, print acceleration, extrusion volume multipliers, print height from reactor base, and suspension viscosity (data not shown). To aid with replication and advancement of our system, all GCODE used for printing has been hosted on a public GitHub repository (https://github.com/tmolley2/Vascular-printing.git). For all vascular printing, a 0.41μm diameter syringe tip was used (22G Nordson EFD tip) given it is within the standard range for mimetic vasculature literature^30,31^. During optimization, we noticed that when the Pluronic was printed at a rate of 5mm per second or higher, the ink tended to over-extrude causing the print to break apart (**Figure 3B**). We found that 2.5 mm/sec gave the most consistent channel width when printing, so we proceeded with this extrusion rate for our vasculature printing used for cell studies. However, when printing high curvature regions, such as those depicted in Figure 1D, a speed of only 0.33mm/second was best for maintaining overall print shape. More printing guidelines can be found in the supplemental methods. By printing the ink back over itself, separately printed channels can be joined to create hierarchical architectures (**Figure 3C**) (Supplemental Video 1). Finally, to establish the broad potential for freeform vascular printing of complex paths, we printed a large perfusable spiral construct in a 5mL suspension (**Figure 3D**) (Supplemental Video 2).

Given that printing resolution is inherently limited by the size of microparticles in a suspension^16^, we wanted to verify that our channels maintained similar topology and wall stress along their entire length. We additionally wanted to verify if the flow characteristics were comparable to native blood vessels. To accomplish this, we performed MicroCT on printed single and bifurcated channels to create a 3D model of the void space. The microgel’s high protein and water content allowed us to segment the microgel volume against air in the channel rather than using contrasting agents. Computational fluid dynamics (CFD) analysis was performed on the segmented channel volumes to measure wall shear stress along the channel lengths under theoretical flow (150nL per second for straight channel, and 300 nL per second for the bifurcation) (**Figure 3E**). The variation of shear stress along the channel varies by ∼100% while also achieving a similar stress level to the theoretical/ideal channel design. Given that blood vessels experience a stress range from 3-30 dyn/cm^2^, we find this variation to be acceptable^31^. Fluid flow vectors also show the fluid path in both channel types (Figures S8-9). Laminar flow is seen without the presence of eddies in both channel types, with similar flow patterns between the experimental and the theoretical channel designs (supplemental videos 3-6).

Having established a method to fabricate hollow channels within the microgels, we next investigated the ability to integrate prototype vascular cells. Human umbilical vascular endothelial cells (HUVECs) were injected into the channels (10 million cells/ml) to create vascular linings. The gels were rotated every 30 minutes for 1 hour to allow the cells to attach to the luminal surface on both sides. After 5 days of culture the HUVECs were seen to adhere and proliferate to the undulating topology of the channel with clear vessel linings at the luminal surface (Figure 4A and B). However, gaps remain in regions between microparticles making the linings incomplete (Figure 4C), and work is ongoing to establish tight linings between all cells. At this stage, we can define blood vessel-like structures within a microgel matrix containing dispersed stromal cells.

**Figure 4.**
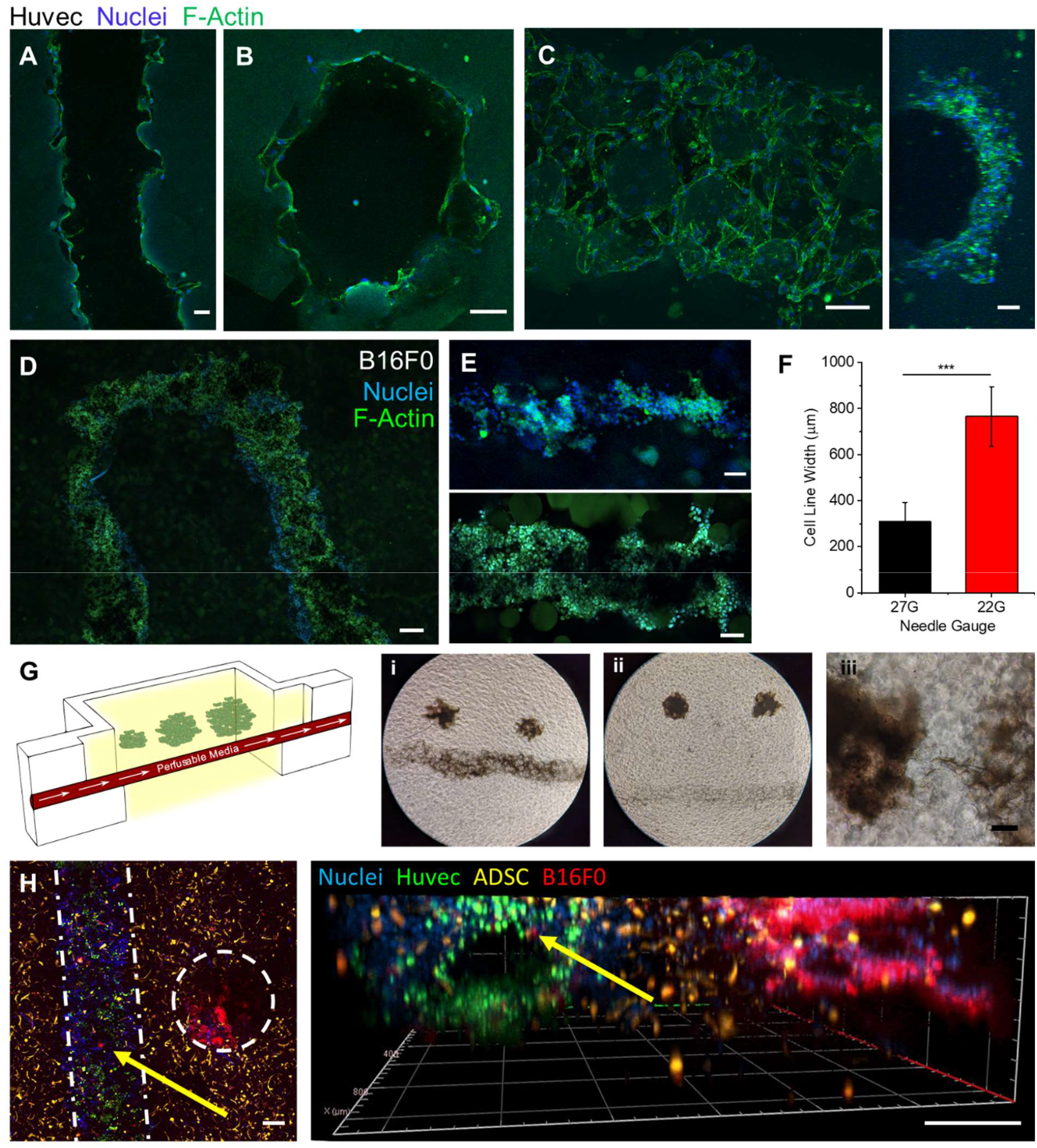
Vascular cell seeding and cell printing. a) Confocal plane of Huvec cells seeded along the walls of a printed channel after 5 days. b) A cross section confocal image of the same gel to verify endothelial cells along the entire channel circumference. c) A max intensity z-stack projection (left, ImageJ) of the top half of a channel of endothelial cells after 5 days (Huvecs) along with a 3D projection image of the side view of that channel (right). d) Confocal image of the top (z-plane) of a U print of a fluidized cell pellet (B16F0). e) Confocal images of printed tumor lines (B16F0) from a 27G needle (Top) and 22G needle (Bottom). f) Plot of the measured average width of tumor line prints from 22G and 27G needles (n=6, P<0.001). g) A schematic of the tumor invasion model (left) and phase contrast images of tumors printed close (i) and far (ii) from the vasculature. A 10x phase contrast image of tumor cells migrating towards vasculature in a close print (iii). h) Confocal Z-projection of a triculture of ADSCs, Huvecs in a channel, and printed B16F0 cell pellets (right) along with a 3D projection (Zen blue, Zeiss) of that same gel (left). Scale bars: 100µm (a, b, c, e, g), 200μm (d, h)

To investigate the propensity for including tumor-like structures, a B16 mouse melanoma tumor model was selected due to its high invasiveness and characteristic black color from melanin production which aids with visualization^32^. A cell pellet fluidized with a 3:5 ratio of culture medium to cells was chosen as the cell ink for simplicity and a high cell density for in situ spheroid production. Parallel lines of printed tumor tissue were first fabricated for viability assessment through live/dead imaging (Figure S10). The cell ink was readily extruded and maintaining its form while printing with little leakage into the void space between microparticles (**Figure 4D**). Cancer cell line thicknesses can be readily controlled by varying the diameter of the nozzle tip used (**Figures 4E-F**). For further modularity, complex shapes can be printed as well as fused together such as rings and thick discs of tumor (Figure S11).

As a proof of concept to demonstrate the capabilities of this platform, we combined tumor printing with vascular printing to create a novel tumor invasion model. We began with printing B16 tumor droplets at distances of either 1mm or 3mm from the vascular channel (**Figure 4G**). Strikingly, before tumor-mediated angiogenesis begin, the cancer cells within 1 mm distance invaded the vasculature in under 4 days (**Figure 4G(i-iii)**). The tumor droplets at 3mm distance showed little invasion. As paracrine signal strength greatly determines cell response^33^, we hypothesize that these tumors may be too far from the vascular lining to interact with it. As further demonstration of the platform’s modularity, a triculture model was created by performing both vascular printing and tumor printing simultaneously in a microgel bath laden with ADSCs throughout the matrix. Here, all three cell types can be seen segregated into their desired locations (**Figure 4H**). After five days, tumor cells tagged with cell tracker deep red can be seen intravasating from the tumor mass into the vasculature (Figure 4H, yellow arrows; Figure S12).

## Discussion

The advent of freeform bioprinting five years ago has led to a rapid development in tools to reconstruct tissue-like structures for model development and tissue engineering applications^13,15^. Recent work by Feinberg and colleagues and Angelini and colleagues has provided new avenues for bioprinting that obviated the need for overly viscous inks through the use of yield stress fluid support baths, enabling 3D printing of intricate structures with broad flexibility in materials selection^14,16^. The two main suspension bath materials used with this printing technique include Carbopol and gelatin microspheres. While these suspensions have excellent yield stress fluid characteristics that allow for ease of printing, they are typically removed post print.

Rather than using the microgel suspension as a sacrificial printing medium, here we recognized the potential for a new class of spatially addressable extracellular matrix, in which cellular activity may be dictated by the properties of the suspension. The Segura group and others have explored these types of granular gels as a cell seeded scaffold to capture the benefits the porous nature of the scaffolds provides^34–36^, thereby demonstrating the potential for cells to be integrated with microgels. In an interesting twist to the composition of the yield stress fluid for printing, Lewis and colleagues demonstrated freeform vascular printing in a suspension of pure cell organoids^23^. Recently, Patrício et al. showed freeform printing of a sacrificial ink into a alginate microgel bath^37^. Following on from these studies, we leveraged the benefits associated with the microporous nature of the support bath to create a cell-laden tunable bioactive matrix, where multiple cell types can be spatially integrated. To demonstrate this, we printed a sacrificial Pluronic-based ink^20^ into cell-microgel suspensions. Coupled with a photocrosslinkable filler polymer between the individual microgels, we stabilized the gels post print and removed of the Pluronic ink. In this way, we constructed complex channels within a cell-laden matrix, that were further modified with endothelial cells towards well defined prototype vessels.

A major advantage with printing in granular media is the ability to print very low viscosity inks without the need for an ink drop printer. Alsberg and colleagues demonstrated this by printing pure pellets of stem cells into their alginate particle baths^17^. In a similar way, we printed tumor aggregates of varied shapes and sizes. Importantly, our approach is the first example of a platform where cellular aggregates can be spatially defined in the presence of uniformly dispersed cells and interspersed vascular channels. We demonstrated this by printing microtumors of melanoma cells at varying distances from prototype vessels and demonstrated distance-invasion relationships. Microfluidic systems with adjacent chambers and counterflow arrangements have been demonstrated to serve as complex heterotypic models to monitor signaling between multiple cell types^38^. However, these platforms invariably involve cells adherent to 2D surfaces which disallows variation in the biochemical and biophysical properties of the microenvironment. Our printing system allows similar associations to be fabricated and monitored in a single bioreactor, in a 3D context with tailorable chemistry and mechanics, thereby providing a more biomimetic environment to study cellular processes.

In conclusion, we have demonstrated a new bioprinting methodology, where a suspension of cells intermixed with chemically stabilized microgels facilitate freeform printing of vascular channels and cellular aggregates in a single chamber. Inspired by the tumor microenvironment, we demonstrate the versatility of this system by integrating prototype tumors and vasculature amidst a matrix of stromal cells. In this way complex processes like tumor intravasation and extravasation, and accompanying roles of stroma-cancer cell interaction, can be readily modelled. Coupled with the ability to simultaneously deposit additional cells with a high degree of spatial control, virtually any number of cell types may be integrated. This new 3D coculture method may provide a means to investigate not only cancer and disease modeling but understanding the role of the extracellular matrix on other cellular processes including tissue morphogenesis in development and disease. Moreover, the high throughput nature of 3D printing combined with this modular approach will allow for combinatorial drug studies to be performed in well-defined physiologically relevant models.

## Materials and methods

### GelMa synthesis

GelMa was synthesized as previously described^3926^. Briefly, gelatin from porcine skin, Type A (Bloom strength 300, Sigma-Aldrich) was dissolved at 10% (w/v) in 1X phosphate buffered saline (PBS, pH 7.4) under stirring at 50°C. 5% (v/w) methacrylic anhydride (Sigma-Aldrich) was added and the mixture stirred for 90 minutes. The solution was diluted two-fold with 1X PBS and centrifuged (3000 rcf, 3 minutes) to remove unreacted methacrylic anhydride particulates. Following this, it was transferred into 14kDa cutoff cellulose dialysis tubes and dialyzed at 40°C for 5-7 days against deionized water. The dialyzed solution was lyophilized for 5-7 days and the resulting powder stored was stored at - 20°C.

### GelMa Microparticle Synthesis

The GelMa microparticles were prepared using a modified water in oil emulsion method^40^. The lyophilized GelMa was hydrated to a 10% (w/v) volume solution in 1X PBS at 40°C. The solution was added dropwise through a 0.45μm sterile filter into a continuously stirring bath of oil (Canola, Sunflower, Olive) (Community co., IGA Australia; Bertolli) at 40°C and allowed to equilibrate for 10 minutes. The bath was cooled to 10°C for 30 minutes prior to adding acetone (22mL/mL GelMa) to dehydrate the microparticles. The particles were then allowed to settle to the bottom of the vessel, washed thoroughly with acetone, and sonicated to break up aggregates. Unbroken aggregates were removed by filtration. The dehydrated microparticles were stored in acetone until use. For size characterization, particles were rehydrated in DI water for one day before taking images on a phase contrast microscope. 100 particles were imaged and their diameters were calculated using ImageJ.

To prepare the microparticles for printing, acetone was removed by evaporation. The microparticles were hydrated for at least 24 hours in a 1% (w/v) solution of GelMa and 0.05 wt% Lithium phenyl- 2,4,6-trimethylbenzoylphosphinate (LAP Sigma-Aldrich, 900889) in either PBS, or appropriate cell culture medium, to achieve a packing fraction of 30% and a final concentration of 1 wt% GelMa in the filler phase as these were determined to be optimal conditions for printing.

### Swell study

A 10wt% solution of pure GelMa dissolved in 1xPBS was warmed in an incubator at 37°C until fully melted. The gel solution (80 µL) was subsequently added to 6×6×2.5mm plastic PLA molds and left at room temp to physically crosslink. Once crosslinked, the gels were weighed and placed into 15mL falcon tubes where they were covered with Acetone (10mL, Chem-supply) and left to shake for 24 hours. The acetone was then decanted, and the gels were air dried for 24 hours to remove all remaining acetone. The dried gels were then weighed before placing into tubes filled with DI water at room temperature. At each time point, the gels were taken from the tube and the surface water was removed with a Kimwipe prior to weighing. The swelling ratio was calculated using the following where W is the weight:

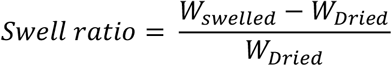

### Rheology

All rheological measurements were performed on an Anton Paar MCR 302 Rheometer with a parallel plate geometry (25mm Disk, 1mm measuring distance, 600uL of suspension bath or Pluronic gel). Oscillatory measurements were performed with 0.02% strain and a 1 Hz frequency for the duration of gelation at 20°C. For in situ UV crosslinking for the GelMa baths, a UV light (with 395nm UV light at 40mW/cm2 for 60 seconds) was placed underneath to illuminate the sample through the quartz crystal stage. Shear rate sweeps were performed with a 1 Hz frequency from a 0.01 to 10 shear rate (1/s) at a log ramp scale over 4 minutes. Temperature stability studies for GelMa baths were run with a temperature ramp from 20°C to 37°C. For the melted samples test, the gels were first placed in an incubator at 37°C for one hour before placing on the rheometer and cooling down to 20°C before running the test. Strain sweep test were performed with a log ramp up rate from 0.02% shear strain up to 200% at 1 Hz frequency over 8 minutes. For the Pluronic temperature sweep, the samples were cooled down in the fridge to 4°C before placing them on the rheometer at 1°C. The temperature was ramped up from 1°C to 37°C at a rate of 1°C/minute with a 0.02% shear strain at 1 Hz frequency. The frequency sweep was run with a log ramp up rate from 0.01 to 100 Hz with a 0.02% strain.

### Atomic force microscopy (AFM)

Suspensions of 30Vf GelMa particles with filler were crosslinked in 6×6×1mm plastic molds glued down to glass coverslips. Shorter molds were used to limit light diffraction for the camera on the AFM’s microscope. The samples were fixed to the bottom of fluorodishes (Coherent, FD35) with 2 part rubber cement. The samples were then submerged in water until ready. All data was acquired with the JPK NanoWizard4 Bio-AFM with a spherical probe (2 µm diameter Borosilicate unmodified probe, Novascan). The tip spring constant was calibrated on glass in water prior to the experiment. Using contact-force microcopy mode, 36 force curves (6 µm approach at 0.5 µm per second) were taken per 10×10µm regions in different locations of the gel. A stitched optical image was taken to find particles and filler spaces between. The curves were loaded in the JPK Data Processing software to calculate the elastic modulus at each region. The following analysis steps were performed:

1. Gaussian smoothing of the curve with a smoothing width of 3.00
2. Baseline subtraction with tilt using the last 40% of the curve along the x-axis
3. Automatic contact point adjustment
4. Vertical tip position calibration using the unsmoothed height.
5. An elasticity fit using the Hertz/Sneddon model with a spherical tip shape with a 1μm tip radius and 0.50 Poisson ratio

### Cell culture and seeding in bulk suspensions

The B16F0 (ATCC) cells were cultured with high glucose Dulbecco’s Modified Eagle Medium (DMEM) supplemented with 10% FBS and 1% Penicillin/Streptomycin. Adipose derived stem cells (ADSCs, PSC-500-011 ATCC) were cultured in low glucose DMEM supplemented with 10% FBS and 1% Penicillin/Streptomycin. GFP-WM266-4 cells with a KO of Elkin1 clone (a gift from K. Poole) were cultured in Minimum Essential Medium Eagle (MEME) with 10% FBS, 1% Glutamax, and 1% Penicillin/Streptomycin. HUVECs (Lonza C2519A) were cultured with the Endothelial Cell Growth Medium-2 BulletKit (Lonza CC-3162) All cultures were maintained at 37°C, 5% CO2 and used between passages 2-13. For ADSCs seeding in the hydrogel matrices, the cells were detached with trypsin, counted, centrifuged down, and resuspended to 2 × 10^7^ cells/mL. The cells were then added in a 1:20 volume ratio to either a solution of 10 wt% GelMa at 37°C or a prehydrated bath of GelMa particles at room temperature for a final concentration of one million cells/mL. These solutions were subsequently supplemented to 0.05wt% LAP with a 2.5 wt% stock. 80uL of each gel solution was then added to plastic printed molds (6×6×2.5mm) where they were crosslinked under a 395nm light torch (Ebay; *100 LED 395 nm UV Ultra Violet Flashlight Blacklight Torch*) at 40mW/cm^2 for 60 seconds. The cells were added to a 24 well plate with 1ml of media. Media was changed after one day followed by every other day. The gels were cultured for 1-7 days before fixation with PFA.

### Cell viability analysis

For ADSCs, 1 million cells per ml were loaded into both the 10 wt% GelMa solution and 30-volume fraction microgel bath, each with 0.05 wt% LAP. Next, 80uL of gel was placed into a 6×6×2.5mm plastic mold where the gels were crosslinked for 1 minute. For the B16F0s, the cell ink was prepped as specified elsewhere. Three lines (22G needle, 5mm long) of cancer cells were printed into each gel prior to crosslinking. All cell loaded gels were placed into a 24 well plate and cultured for the specified time. Media changes were made on days 1, 3, and 5. For the staining, the media was removed and the gels were washed once with PBS prior to the addition of 500uL of 1X PBS with Calcein AM (2 µM) and Ethidium Homodimer-1(4 µM) (Invitrogen, L3224). After 45 minutes of incubating the stains, the gels were rinsed with PBS and washed again with PBS after 10 minutes before imaging on a Zeiss LSM 800 Confocal microscope.

### Immunofluorescence staining and tissue clearing

Clearing solutions were prepared as done previously with minor modifications (Molley 2020, susaki 2014). Briefly, Cubic solution 1 was prepared by mixing 25 wt% urea (Sigma Aldrich., 583051), 25 wt% N,N,N’,N’-tetrakis(2-hydroxypropyl) ethylenediamine (Sigma Aldrich, 585714), and 5 wt% Triton X-100 (Sigma Aldrich, 562380) into DI water at 50°C until fully dissolved. Cubic solution 2 was prepared by mixing 50 wt% sucrose (Sigma Aldrich, 584173), 25 wt% urea, 10 wt% triethanolamine (Sigma Aldrich, 90278-100mL) with DI water at 55°C until also fully dissolved. Microgel suspensions were fixed using a 4 wt% paraformaldehyde (Chem-Supply) for 1-4 days at room temperature to ensure fully penetration of PFA into thick constructs. The gels were then rinsed with PBS followed by 3 PBS washes at 2-4 hour intervals. Gels that required antibody staining were then placed into a 5ml Eppendorf tube filled with Cubic clearing solution 1 for 48-96 hours. The gels were then rinsed with PBS followed by 3 PBS washes at 2-6 hour intervals. Primary antibody stains were diluted in 1x PBS with 1 wt% Bovine Serum Albumin (BSA) and added to the gels for 24 hours at room temperature. The gels were then rinsed with 1x PBS (with 1wt% BSA) followed by 3 PBS (with 1 wt% BSA) washes at 2-6 hour intervals. The secondary antibody staining was then performed in 1% BSA solution in PBS with Hoechst and 488-Phalloidin. The gels were washed with PBS three final times before the addition of the Cubic 2 clearing solution for 2-5 days. All confocal imaging was performed with a Zeiss LSM 800. A 10x objective with a 2.5mm working distance was used to see deeper into the samples. Samples were coated with clearing 2 solution throughout the duration for the imaging to prevent drying.

### Cell volume segmentation analysis

For cell volume analysis, one million ADSCs were loaded into microgel suspensions and bulk hydrogels before crosslinking for 60 seconds. At the desired time points, the cells were fixed with 4% PFA for 24 hours before staining (Hoechst, 405; Phalloidin, 488) and cleared as mentioned above. Confocal z-stacks (20x objective, 109 slices over 50microns) were taken of representative regions in each gel. The images were imported in Imaris for analysis. Cell segmentations were created with identical thresholding values per gel with each independent nucleus as a seed for the cells.

### Plastic reactor mold fabrication

All plastic reactor molds were 3D printed with a Lulzbot Mini2 plastic 3D printer with a 0.25mm nozzle end. For cell experiments, molds are fixed to an 18mm diameter glass coverslip with cyanoacrylate glue. The molds are then quickly soaked with 80 vol% ethanol and dried out inside of a biosafety cabinet prior to use. For non-cell experiments, the reactors are pressed into stretched parafilm before addition of the microgel suspension and subsequent crosslinking. STL files for the molds can be found here: https://www.thingiverse.com/tmolley/collections/fre eform-vascular-printing-designs

### Pluronic ink preparation

To create the sacrificial inks, Pluronic F127 (Sigma, P2443-250G) was first weighed out into 50mL flacon tubes. Cold DI water (4°C) was then added to the Pluronic powder for the appropriate weight percentage. The mixture was mechanically agitated before placing into a fridge at 4°C overnight to fully dissolve the ink. The ink was then stored at 4°C until further use.

### MicroCT and Volume segmentation

MicroCT scan was performed with the U-CT (MILabs, Utrech), with 50 kVp x-ray tube voltage, 0.21 mA tube current, 75 ms per frame, 360° angle, and 0.25° projections. Images were reconstructed with MILabs Recon 10.16 at 20 µm voxel size and vessels segmented using Imalytics Preclinical 2.1 (Gremse-IT GmbH, Germany).

### CFD and analysis

Computational fluid dynamics (CFD) analysis was run with the Autodesk CFD 2019® software. The segmented STL meshes exported from Imalytics Preclinical 2.1 were imported in Autodesk 360 Fusion® to reduce the mesh network down to <10,000 polygons for smoother modeling. Theoretical model designs were created in Autodesk Inventor CAD to represent the shape the gcode was supposed to create. The mesh volumes were the loaded in the CFD software and the following assumptions were made:

1. Volume is specified as water
2. End boundary condition set to 0 Pa pressure
3. Automatic meshing
4. 0 initial conditions
5. Fluid is incompressible
6. Flow was set to a kappa-epsilon turbulent flow model with a turbulent:lamilar flow ratio of 100:1
7. ADV 5 modified Petrov-Galerkin Advection
8. 100 interations were performed with a steady state solution mode
9. Flow rate defined as 150nL/s for the straight channel and 300nL/s for the bifurcation
10. The bifurcation had flow originating from the single channel end

Videos of flow traces were recorded and exported from the software.

### Printing (vasculature, tumors, co-culture cell baths)

#### Printing Vasculature

A Lulzbot mini2 retrofitted with a screw extrusion syringe head (Replistruder head 2, Feinberg lab) was placed into a Biosafety cabinet. For Pluronic printing, the 29wt% Pluronic F127 solution was cooled down in a fridge (4°C), then pulled into an airtight glass syringe (Hamilton® 1002LTN syringe) and inverted to remove air bubbles. The syringe was warmed to room temperature to gel the Pluronic F127 before loading into the printer. A 22G Nordson EFD needle tip was added to the syringe and a small amount of Pluronic was extruded out to prime the needle tip. The print needle was then orientated over and aligned with the inlet and the suspension was added to the mold until the surface of the liquid was flush with the top of the mold. The desired print code was run, and the needle was gently cleaned with a Kimwipe prior to the next print. The suspension was then photocrosslinked for 1 minute, placed into a 12 well plate, parafilmed, and put in a fridge for 15 minutes to liquify the Pluronic F127. For print fidelity measurements, the ink was removed and Phase contrast images of the air inside the channel were taken. Analysis was performed via ImageJ along 6 diameters for each line to determine the lines average thickness.

#### Direct printing of cells

The desired cells were treated with trypsin, centrifuged, washed, and then pelleted. The cell pellets were lightly fluidized with media in a 5:3-5:2 ratio of cells to media to break up aggregates. Care was taken to limit the introduction of air bubbles during this stage. The pellet was then pulled into a 1mL syringe (Livingston), and the syringe was loaded directly into a 3D printed fitting on the bioprinter. The desired syringe needle was then primed with cell solution and printed into molds filled with a microgel suspension.

#### Dual Cell and Vascular printing

Each part of the multistage printing process was performed as mention above with some modifications. Importantly, cell printing preceded vascular printing as the Pluronic ink begins to diffuse into the surrounding suspension if not crosslinked fast enough leading to poor channel resolution. In addition, after cell printing, the molds are placed into a covered, sterile petri dish to enable easy access while limiting overhead airflow that can dry out or contaminate the microgel suspension.

#### Incorporating cells into suspensions

When incorporating cells into the support suspensions, the microparticles were first hydrated with the appropriate culture medium. The cells were treated with trypsin, centrifuged to a pellet, then resuspended to a 50x cells mL concentration compared to the final volume. The high concentration cell solution was gently mixed into the hydrated microparticles before adding to the molds. Printing was then conducted as mentioned prior.

### Loading vascular cells in printed vasculature and subsequent co-culture

The microgels were placed into a fridge for 15 minutes to allow the Pluronic F127 to transition into a liquid state. It was then removed via holes at either end of the mold, leaving behind a hollow channel. Endothelial cells (HUVECs at 10-20 million cells/mL) were loaded into a 1mL syringe and injected into the channel through the same holes at either end of the mold. The microgel was inverted and placed in a 12-well plate, then placed in the incubator for 30 minutes. The vessels were then flipped back upright and incubated for another 30 minutes before adding the cell media. The construct was cultured at 37°C for 4- 7 days.

### Fidelity of tumor prints

For tumor line prints, a sacrificial print was first made above the microgel suspension to prime the needle tip. Once printed, the microgels were immediately fixed with 4% PFA. After fixation and washing of the fixed microgels, they were added to a 5 wt% solution of Hydrogen Peroxide (Sigma Aldrich, 487568) at room temperature for 24 hours to bleach the melanin and aid in confocal imaging. The samples were then stained with Hoechst and Phalloidin and z-stack tile scans of the gels were taken. Analysis was performed in ImageJ. First, the z-stacks were projected into one slice with using the maximum brightness. The images were then thresholded in the phalloidin channel to outline the lines, followed by 6 length measurements taken across the length of the tumor lines.

### NMR For GelMa Methacrylation Characterization

The degree of functionalization (DOF) was quantified using a 1H NMR spectrometer (Bruker Avance III 400 MHz) by referencing 1H NMR chemical shifts to the residual solvent peak at 4.80 ppm in D2O. Briefly, 10 mg of GelMA was dissolved in 1 mL of D2O at 37°C. 700 µL was put into an NMR tube for the acquisition of the NMR data. NMR spectra were analyzed using MestReNova (Mestrelab Research) by Dr. Julio Serrano (University of Illinois at Urbana-Champaign), using the chemical shift in the aromatic region as integral reference. Degree of functionalization of 96 and 95% can be seen in Figure S13. 1 H NMR (400 MHz, D2O,): δ 7.24 (m), 5.65 (m), 5.40 (m).

### Statistical analysis

The whiskers in the box plots are standard deviation (s.d.) unless otherwise specified. Statistical significance was determined using a one-way ANOVA with Tukey’s Post Hoc HSD analysis. Differences were considered significant when P < 0.05.

## Acknowledgments

This work was supported through funding from the National Health and Medical Research Council Grant # APP1185021. This material is also based upon work supported by the National Science Foundation Graduate Research Fellowship Program and the National Science Foundation Graduate Research Opportunities Worldwide program under Grant No. DGE- 1144245 (A.S.T.). We acknowledge the help and support of staff at the Biomedical Imaging Facility and the Biological Specimen Preparation Laboratory of the UNSW Mark Wainwright Analytical Centre. The authors declare no conflicts of interest.

## References

1. Malandrino, A., Kamm, R. D. & Moeendarbary, E. In Vitro Modeling of Mechanics in Cancer Metastasis. ACS Biomater. Sci. Eng. 4, 294–301 (2018).

2. Nicolau, M., Levine, A. J. & Carlsson, G. Topology based data analysis identifies a subgroup of breast cancers with a unique mutational profile and excellent survival. Proc. Natl. Acad. Sci. U. S. A. 108, 7265–7270 (2011).

3. Tschirhart, C. E., Nagpurkar, A. & Whyne, C. M. Effects of tumor location, shape and surface serration on burst fracture risk in the metastatic spine. J. Biomech. 37, 653–660 (2004).

4. Bailey, J. M., Mohr, A. M. & Hollingsworth, M. A. Sonic hedgehog paracrine signaling regulates metastasis and lymphangiogenesis in pancreatic cancer. Oncogene 28, 3513–3525 (2009).

5. Kwon, M. C. et al. Paracrine signaling between tumor subclones of mouse sclc: A critical role of ets transcription factor pea3 in facilitating metastasis. Genes Dev. 29, 1587–1592 (2015).

6. Chen, Y. et al. EMMPRIN regulates tumor growth and metastasis by recruiting bone marrow-derived cells through paracrine signaling of SDF-1 and VEGF. Oncotarget 6, 32575–32585 (2015).

7. Fidler, I. J., Yano, S., Zhang, R. D., Fujimaki, T. & Bucana, C. D. The seed and soil hypothesis: Vascularisation and brain metastases. Lancet Oncology 3, 53–57 (2002).

8. Kenig, S., Alonso, M. B. D., Mueller, M. M. & Lah, T. T. Glioblastoma and endothelial cells cross-talk, mediated by SDF-1, enhances tumour invasion and endothelial proliferation by increasing expression of cathepsins B, S, and MMP-9. Cancer Lett. 289, 53–61 (2010).

9. Rao, S. et al. CXCL12 Mediates Trophic Interactions between Endothelial and Tumor Cells in Glioblastoma. PLoS One 7, e33005 (2012).

10. Upreti, M. et al. Tumor-Endothelial Cell Three-dimensional Spheroids: New Aspects to Enhance Radiation and Drug Therapeutics. Transl. Oncol. 4, 365–376 (2011).

11. Wang, C. et al. Mimicking brain tumor-vasculature microanatomical architecture via co-culture of brain tumor and endothelial cells in 3D hydrogels. Biomaterials 202, 35–44 (2019).

12. Kam, Y., Rejniak, K. A. & Anderson, A. R. Cellular modeling of cancer invasion: Integration of in silico and in vitro approaches. J. Cell. Physiol. 227, 431–438 (2012).

13. Bhattacharjee, T. et al. Writing in the granular gel medium. doi: 10.1126/sciadv.1500655

14. Morley, C. D., Tordoff, J., O’Bryan, C. S., Weiss, R. & Angelini, T. E. 3D aggregation of cells in packed microgel media. Soft Matter 16, 6572–6581 (2020).

15. Hinton, T. J. et al. Three-dimensional printing of complex biological structures by freeform reversible embedding of suspended hydrogels. Sci. Adv. 1, e1500758–e1500758 (2015).

16. Lee, A. et al. 3D bioprinting of collagen to rebuild components of the human heart. Science 365, 482–487 (2019).

17. Jeon, O. et al. Individual cell-only bioink and photocurable supporting medium for 3D printing and generation of engineered tissues with complex geometries. Mater. Horizons (2019). doi: 10.1039/C9MH00375D

18. Noor, N. et al. 3D Printing of Personalized Thick and Perfusable Cardiac Patches and Hearts. Adv. Sci. 6, (2019).

19. Daly, A. C., Riley, L., Segura, T. & Burdick, J. A. Hydrogel microparticles for biomedical applications. Nature Reviews Materials 5, 20–43 (2020).

20. Wu, W., Deconinck, A. & Lewis, J. A. Omnidirectional printing of 3D microvascular networks. Adv. Mater. 23, H178–H183 (2011).

21. Kolesky, D. B., Homan, K. A., Skylar-Scott, M. A. & Lewis, J. A. Three-dimensional bioprinting of thick vascularized tissues. Proc. Natl. Acad. Sci. U. S. A. 113, 3179–84 (2016).

22. Kobayashi, K. et al. On-site fabrication of Bilayered adhesive mesenchymal stromal cell-dressings for the treatment of heart failure. Biomaterials doi: 10.1016/j.biomaterials.2019.04.014

23. Skylar-Scott, M. A. et al. Biomanufacturing of organ-specific tissues with high cellular density and embedded vascular channels. Sci. Adv. 5, eaaw2459 (2019).

24. Wickström, S. A. & Niessen, C. M. Cell adhesion and mechanics as drivers of tissue organization and differentiation: local cues for large scale organization. Current Opinion in Cell Biology 54, 89–97 (2018).

25. Potier, E., Noailly, J. & Ito, K. Directing bone marrow-derived stromal cell function with mechanics. Journal of Biomechanics 43, 807–817 (2010).

26. Susaki, E. A. et al. Advanced CUBIC protocols for whole-brain and whole-body clearing and imaging. Nat. Protoc. 10, 1709–1727 (2015).

27. Molley, T. G. et al. Geometrically Structured Microtumors in 3D Hydrogel Matrices. Adv. Biosyst. 2000056 (2020). doi: 10.1002/adbi.202000056

28. Lin, N. Y. C. et al. Renal reabsorption in 3D vascularized proximal tubule models. Proc. Natl. Acad. Sci. U. S. A. 201815208 (2019). doi: 10.1073/pnas.1815208116

29. Kolesky, D. B. et al. 3D Bioprinting of Vascularized, Heterogeneous Cell-Laden Tissue Constructs. Adv. Mater. 26, 3124–3130 (2014).

30. Rein, J. L. et al. Effect of luminal flow on doming of mpkCCD cells in a 3D perfusable kidney cortical collecting duct model. METHODS CELL Physiol. Mak. Cell Cult. More Physiol. Am J Physiol Cell Physiol 319, 136–147 (2020).

31. Polacheck, W. J. et al. A non-canonical Notch complex regulates adherens junctions and vascular barrier function. Nature 552, 258–262 (2017).

32. Lee, J., Abdeen, A. A., Wycislo, K. L., Fan, T. M. & Kilian, K. A. Interfacial geometry dictates cancer cell tumorigenicity. Nat. Mater. 15, 856–862 (2016).

33. Wang, Y. H. et al. Vascular endothelial cells facilitated HCC invasion and metastasis through the Akt and NF-κB pathways induced by paracrine cytokines. J. Exp. Clin. Cancer Res. 32, 1–11 (2013).

34. Truong, N. F. et al. Microporous annealed particle hydrogel stiffness, void space size, and adhesion properties impact cell proliferation, cell spreading, and gene transfer. Acta Biomater. 94, 160–172 (2019).

35. Griffin, D. R., Weaver, W. M., Scumpia, P. O., Di Carlo, D. & Segura, T. Accelerated wound healing by injectable microporous gel scaffolds assembled from annealed building blocks. Nat. Mater. 14, 737–744 (2015).

36. Kodali, A., Lim, T. C., Leong, D. T. & Tong, Y. W. Cell-Microsphere Constructs Formed with Human Adipose-Derived Stem Cells and Gelatin Microspheres Promotes Stemness, Differentiation, and Controlled Pro-Angiogenic Potential. Macromol. Biosci. 14, 1458–1468 (2014).

37. Patrício, S. G. et al. Freeform 3D printing using a continuous viscoelastic supporting matrix. Biofabrication doi: 10.1088/1758-5090/ab8bc3

38. Rothbauer, M., Zirath, H. & Ertl, P. Recent advances in microfluidic technologies for cell-to-cell interaction studies. Lab on a Chip 18, 249–270 (2018).

39. Loessner, D. et al. Functionalization, preparation and use of cell-laden gelatin methacryloyl-based hydrogels as modular tissue culture platforms. Nat. Protoc. 11, 727–746 (2016).

40. Phromsopha, T. & Baimark, Y. Preparation of Starch/Gelatin Blend Microparticles by a Water-in-Oil Emulsion Method for Controlled Release Drug Delivery. Int. J. Biomater. 2014, 829490 (2014).

