## Supplemental Figures for "Freeform printing of heterotypic tumor models within cell-laden microgel matrices"

[1]School of Materials Science and Engineering, University of New South Wales, Sydney NSW 2052
[2]Dept. Chemical and Biomolecular Engineering, University of Illinois at Urbana-Champaign, Urbana, IL 61801, United States; Carl R. Woese Institute for Genomic Biology, University of Illinois at Urbana-Champaign, Urbana, IL 61801, United States
[3]Biological Resources Imaging Laboratory, Mark Wainwright Analytical Centre, University of New South Wales, Sydney NSW 2052
[4]School of Chemistry, Australian Centre for Nanomedicine, University of New South Wales, Sydney NSW 2052

### Supplementary experimental methods

*3D printing guidelines*

Bioprinting is a complex fabrication scheme whose success is greatly reliant on a variety of printing parameters and bioink materials properties. Introducing a tertiary component – a support bath – can compound this difficulty with a sleuth of new variables to cause failed prints. Given this multivariate landscape, it can be quite a daunting task to troubleshoot and optimize printing. In an effort to make it easier for other researchers to adopt this method, we have created a list below of tips, guidelines, and common issues we have come across while working on this project. This is by no means an exhaustive list, and it also pertains to our specific setup, materials used, and lab environment. However, while these cannot be taken as gospel, we hope they will aid those who wish to print freeform vascular channels or cell pellets into a suspension bath.

***Preparing the suspension baths for printing***

To ensure reliable results when printing, it is critical to prepare the suspensions in the same way every single time. Given the nature of having a bulk phase and a liquid phase, slight changes to the ratio of particles to surrounding filler can have large effects on the suspension's mechanics and printability. A bath which is too viscous, i.e., contains too much liquid phase, will not be able to hold the print in place, causing it to either be dragged with the needle tip, or sink to the bottom of the suspension. Conversely, a bath which is too viscous will not allow the ink to be deposited into it, causing breaks in the print. Here are some general tips:

1. Ensure particles are fully dry before hydrating. They should look like sand and flow like a liquid
2. Allow the particles at least 12 hours to hydrate, overnight is best. The suspension will gain viscosity as the particles slowly hydrate and take up a larger volume fraction
3. When using strong particles, pipette aggressively when hydrating to break up as many dry particle clumps as possible. If some clumps don't get dispersed, then they will remain a clump once hydrated and not be useful for printing
4. Aim to make twice as much suspension bath per experiment as you need, as the suspension will stick to the sides of tubes, pipette tips, etc. and lower your yield.
5. Try adding the LAP to the filler solution used to hydrate the particles, before adding to the particles. This helps ensure even mixing of the photocrosslinker.
6. Once the bath is hydrated, it is best practice to pipette very slow, and remove bath from deep withing the solution to limit the introduction of air bubbles.
7. If many air bubbles are formed, the solution can be centrifuged at 500-1000rcf for 5 minutes to pull them out.

8. Taller tubes work better at limiting air bubble introduction, such as the 15ml conical tubes.
9. When finally placing the solution into the mold, ensure the suspension is level before you begin printing. If it is under filled and bows inside, the print will become distorted and may come out of solution. If it is over filler, the gels will be harder to image as the light will be curved when it enters the sample, like a lens.

***Pluronic printing into suspension baths***

Printing Pluronic can be a bit of a challenge as it will rapidly change its viscoelastic properties with temperature changes. It is also prone to not anchoring in suspension baths if the printing parameter are not optimizing. Here are some guidelines to follow to get the best reproducibility:

1. Regular clean the needle tip every time you print since excess Pluronic on the side of the needle can disrupt how it deposits in the next suspension bath
2. You may need to change the weight percentage of Pluronic that your lab uses depending on the internal temperature of your lab
3. By having the Pluronic touch the plastic of the mold around it helps anchor it down and greatly increases chance of success. Otherwise, the printed Pluronic may just drag along the bath as it is printed rather than staying where it is deposited.
4. Before each print, it is important to prime the needle with Pluronic as the ink can dry out at the needle interface and not print evenly at the start, causing the beginning of your print to not occur. We have this built into our code where we have a small excess pushed out and the printer waits 3 seconds for us to wipe off the excess before it moves into the suspension to start.

#### *Cell pellet printing into suspension baths*

Printing the cell pellets to create droplets is not as difficult. However, the more complex the same and architecture you desire, the more difficult the code writing will become. Here are some guidelines to help get you started:

1. The cell pellet should be fluidized with at least a small amount of cell media. If not, the cells can be quite clumpy and will not print evenly.
2. It is critical to culture a significantly higher number of cells than you need to print. Many cells will be lost when filling the syringe needle, priming the needle for each set of printing, and loading the syringe into the print head. We found that at least a 150uL cell pellet was sufficient to have enough cells for every set of printing we did.
3. We found it best to retract the volume inside the needle while pulling the needle out of the bath in between each droplet or line that we print. If the needle does not leave the bath, the syringe tends to drag excess cells as it moves, leading to poor fidelity.
4. We also found it important to wash the tops of microgel once crosslinked as cell can be pulled onto the top of the gel from the needle as it comes out of the bath. This can be aided with a good bioprinter that can retract well

#### *Combination printing of vasculature and cells*

Here things get quite tricky as there are many moving parts associated with printing multiple cell types in multiple ways while trying to maximize the viability of all cells. We are still learning more and more each time, so here are some challenges we still face and some guidelines on how to best overcome them:

1. Try printing one ink at a time for all of you baths before switching to the next ink and repeating in all baths i.e. print all cells first in all baths, then print the Pluronic into those baths.
    a. Swapping print heads takes the most time and removes the orientation calibration which will add even more time per print
2. One challenge is that the more samples you need to print at once, the longer the suspension sit before being photocrosslinked. The issue is that the suspension can begin to dry out at the top, causing cracks in the top of the bath that make removing the vasculature difficult. Try keeping the suspension covered under a petri dish to limit drying, and a small amount of fresh suspension can be added to the top of the bath before the second print.
3. We found it best to print the cell pellet into the suspension first, followed by the vasculature. When printing the Pluronic first, it would begin to diffuse into the suspension and create poor fidelity channels while also being more difficult to remove once liquified.

### Supplementary Figures

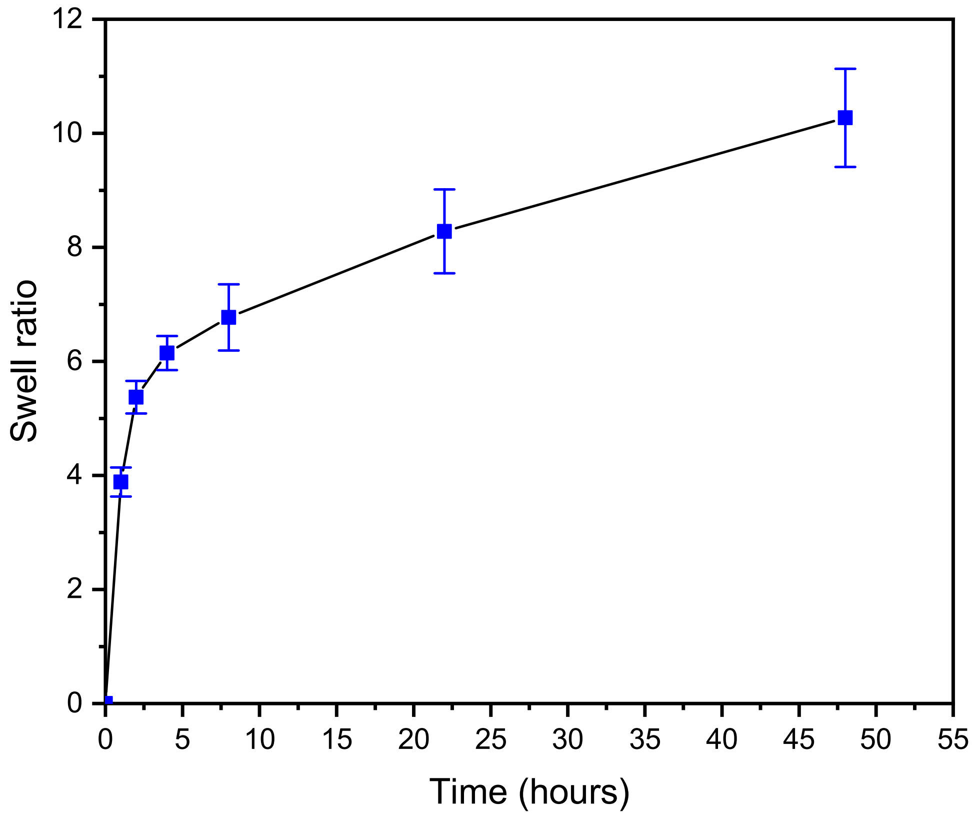


**Figure S1.** Swelling data (n = 5) of pure 10 wt% GelMa hydrogels (75uL) physically crosslinked and dried with acetone to mimic particle drying.

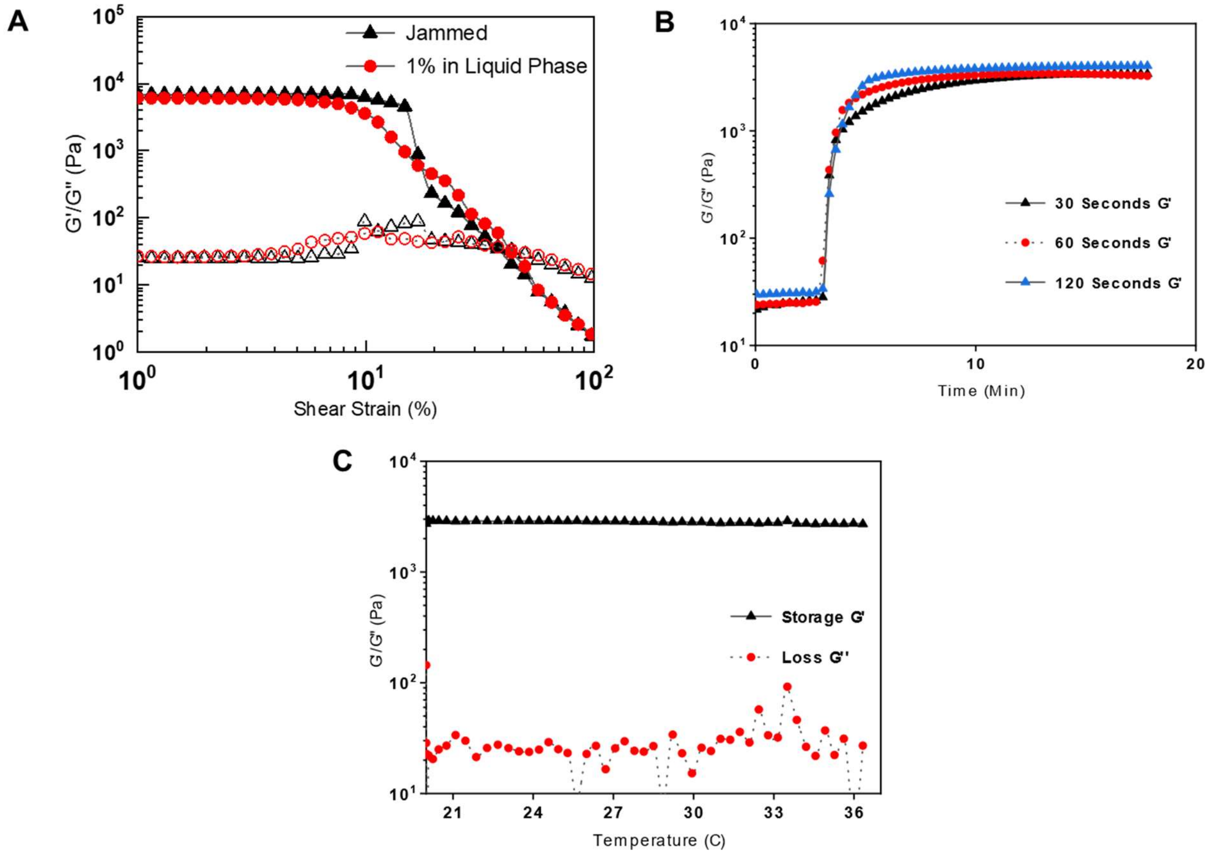


**Figure S2.** Extra Rheology data for microgel suspensions made from 10 wt% GelMa particles. a) Shear strain amplitude sweeps of suspensions with (red markers, 40% volume fraction of particles) and without (black markers, 60% volume fraction of particles) a 1 wt% GelMa filler in the liquid phase. Closed markers are the storage modulus (G') and open markers are the loss modulus (G''). b) Gelation curves of 1% filler GelMa suspensions exposed to 30 (Black Triangle), 60 (Red Circle), and 120 (Blue Triangle) seconds of 395nm light for crosslinking. c) Temperature sweep from 20 to 37°C of a 1% filler GelMa suspension (40% volume fraction of particles)

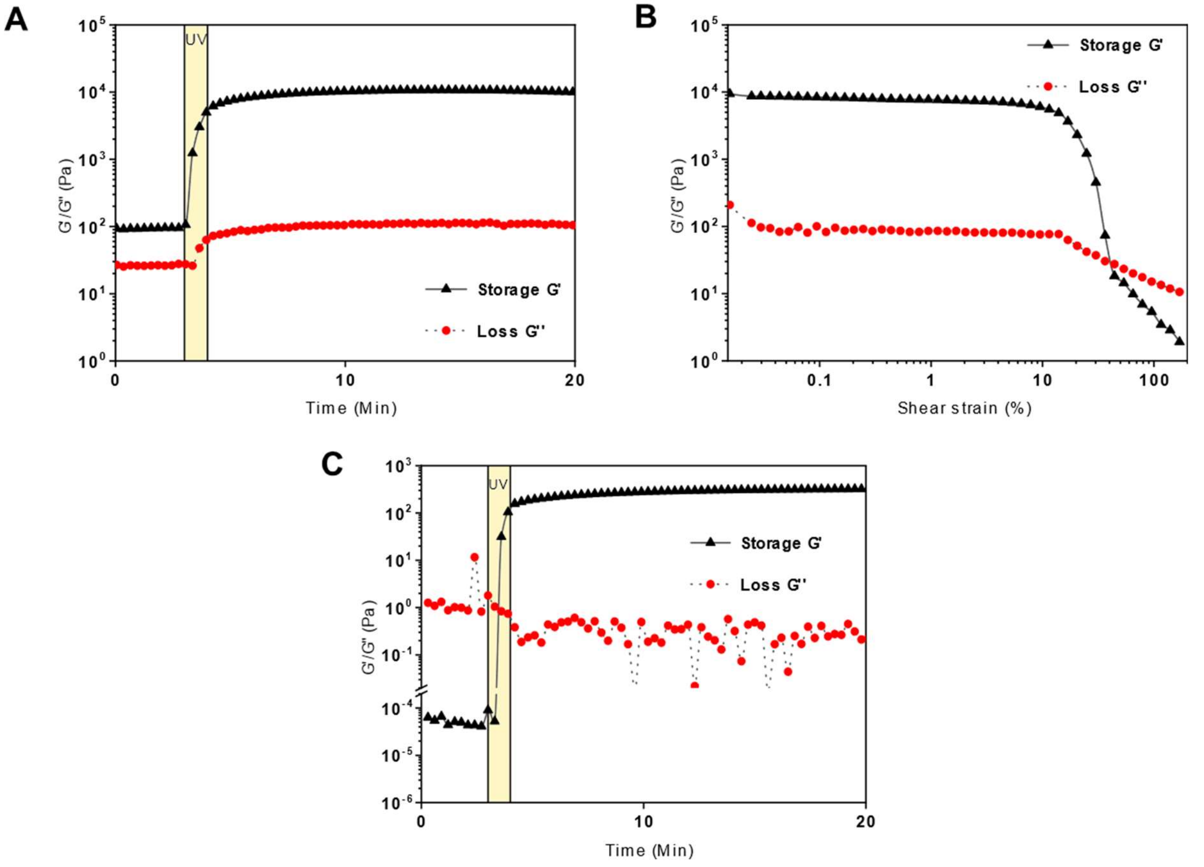


**Figure S3.** Rheology of microgel suspensions made with 15 wt% GelMa Microparticles with 1 wt% GelMa as the filler. a) Gelation curve with 60 seconds of UV. b) Shear strain amplitude sweep post crosslinking. c) Gelation curve of a melted suspensions (37°C for 1 hour to fully melt). For all curves, black triangle markers correspond to the storage modulus (G') and red circle markers correspond to the loss modulus (G").

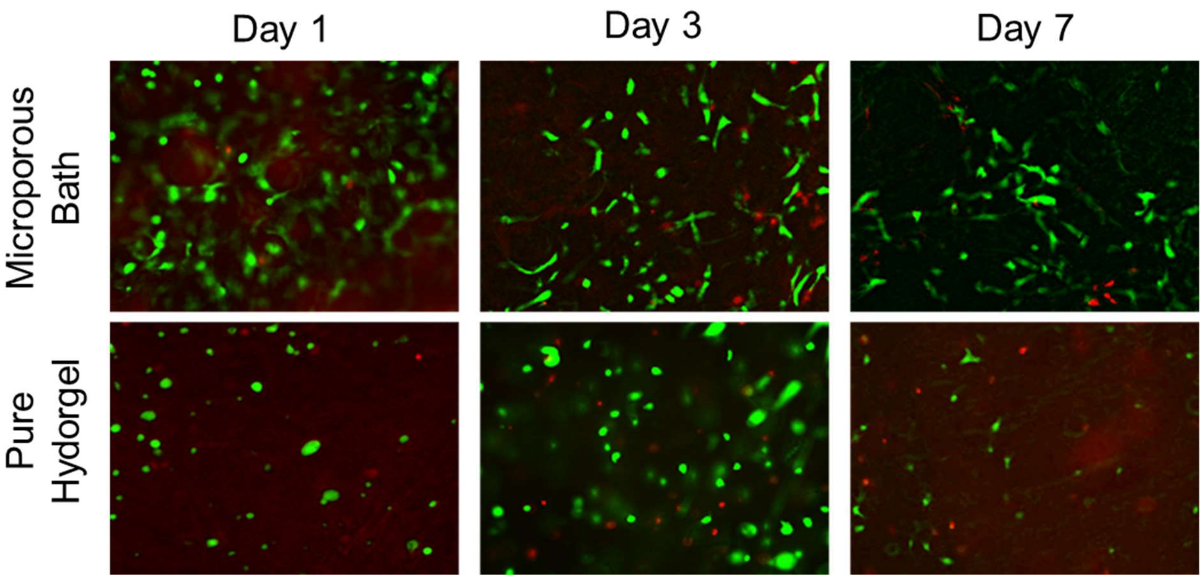


**Figure S4.** Live (green, calcein AM) dead (red, ethidium homodimer-1) stain of ADSCs (1 million cells per ml) loaded into a microgel suspension and pure 10 wt% GelMa hydrogel. Scale bars:

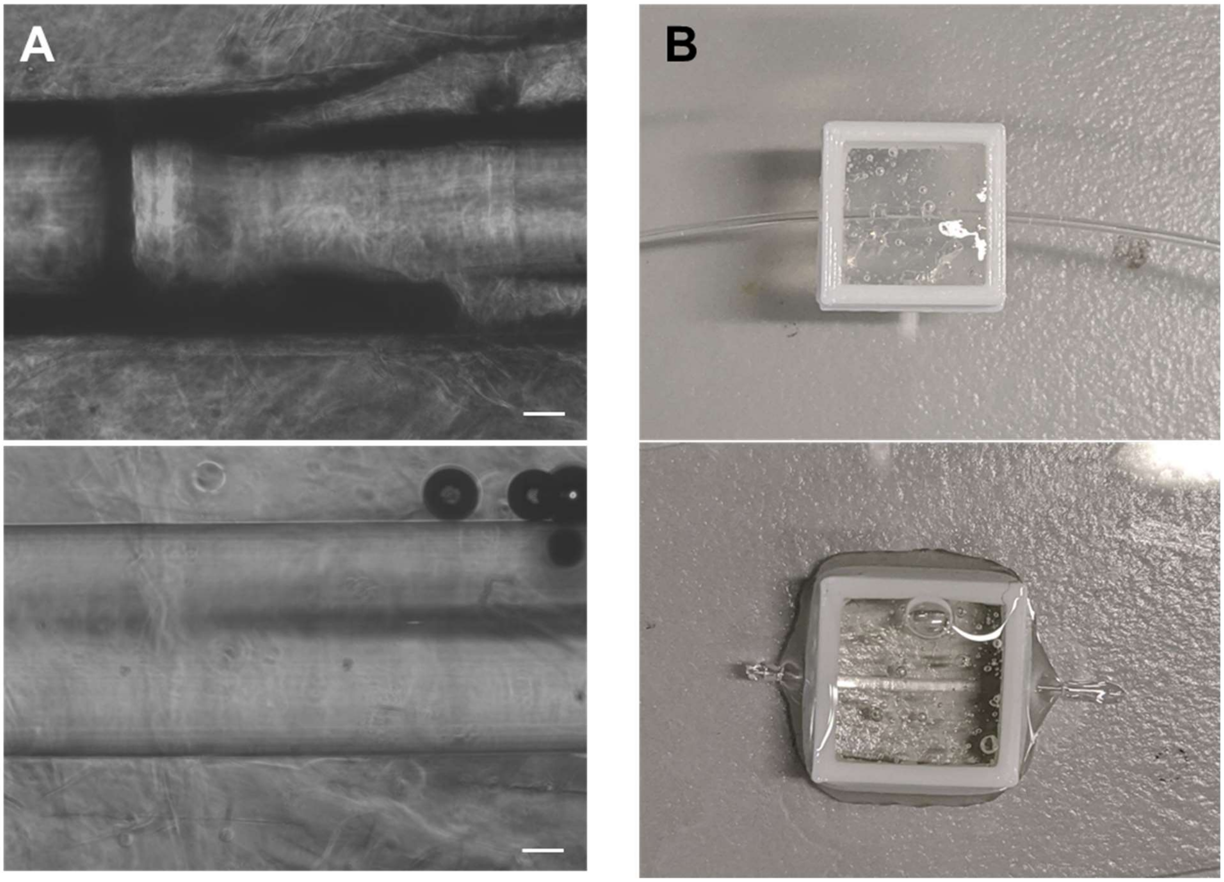


**Figure S5.** Tissue clearing of particle suspensions. Phase contrast images (a) and optical images (b) of a fishing line strewn through a microgel suspension before (top) and after (bottom) adding a tissue clearing solution and incubating for 24 hours. Scale bars: 200µm.

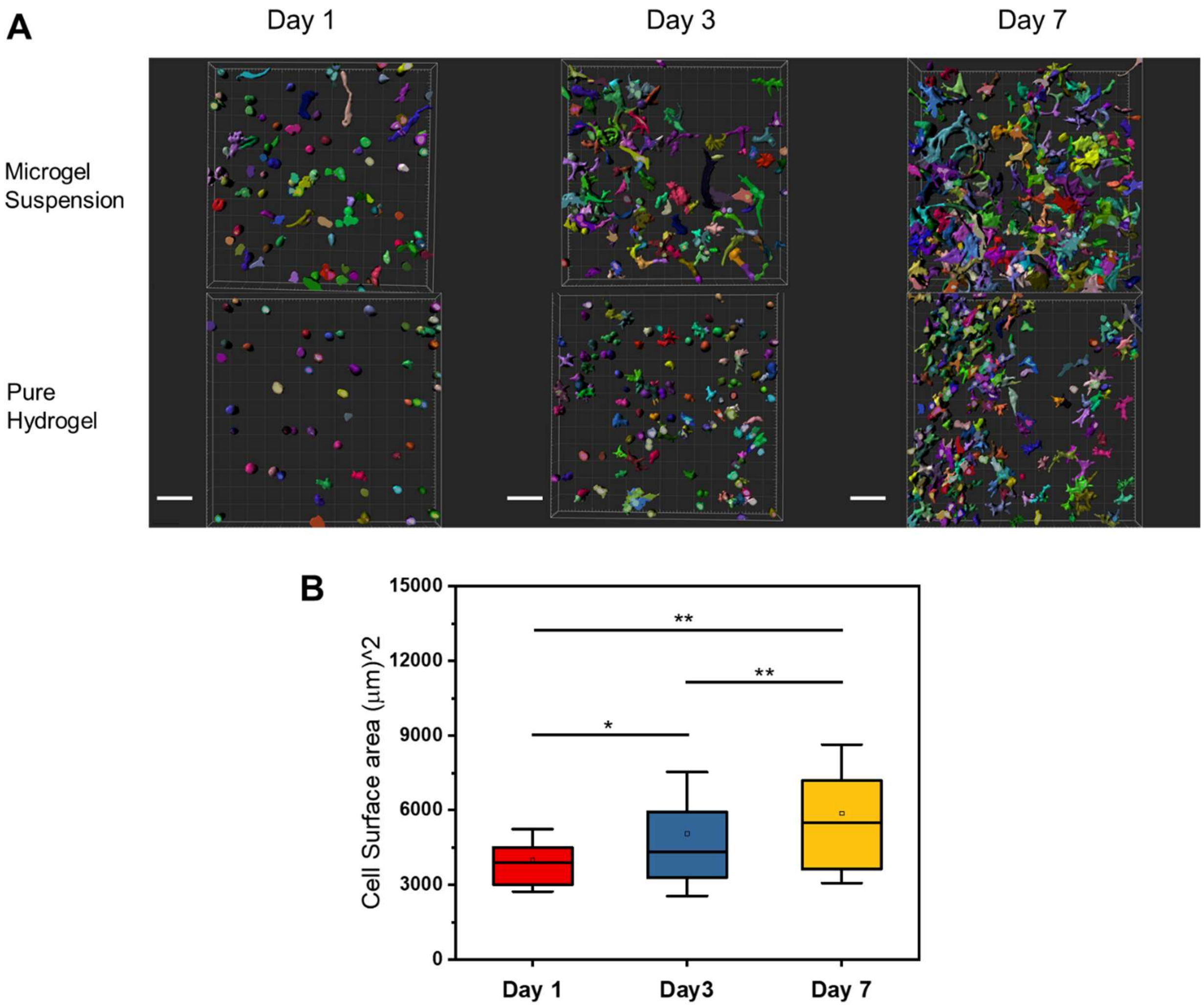


**Figure S6.** Volumetric segmentation of cell volumes. a) 3D segmented volumes of ADSCs from Imaris in Microgel suspensions (Top) and Pure bulk GelMa (Bottom, 10 wt%) at days 1 (left), 3 (center), and 7 (right). b) Cell surface area box and whisker plots for the bulk hydrogel samples. Statistical significance was found between days 1 and 3 ($p = 0.02443$), days 3 and 7 ($p = 0.00914$), and days 1-7 ($p = 0.00101$). Scale bars: 100μm

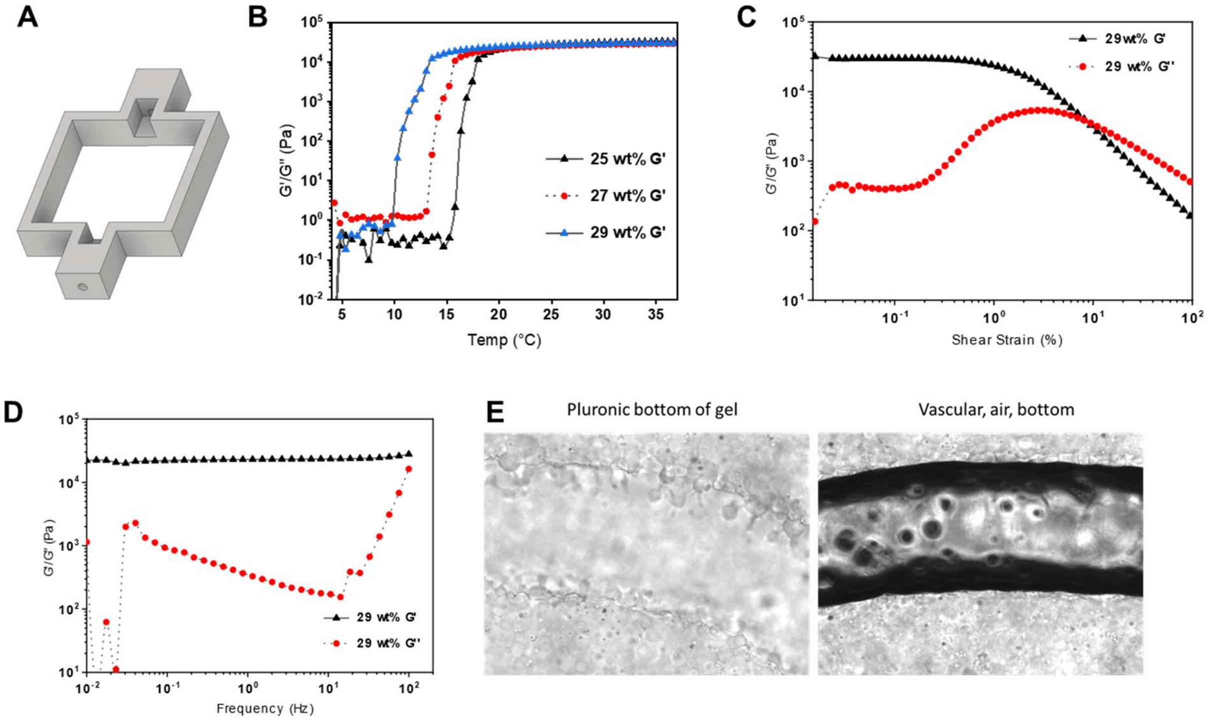


**Figure S7.** Pluronic F127 ink characterization. a) CAD design for the vascular print reactors. b) Rheology temperature ramp (4°C to 37°C at 1.1°C per minute) for the storage modulus of 25 wt% (black markers), 27 wt% (red markers), and 29 wt% (Blue markers) Pluronic F127. c) Shear strain amplitude sweep for the storage (black markers) and loss (red markers) moduli of 29 wt% Pluronic d) Frequency sweep for the storage (black markers) and loss (red markers) moduli of 29 wt% Pluronic e) Phase contract images of Pluronic printed in a microgel (left) followed by the removal of that Pluronic from the microgel (right)

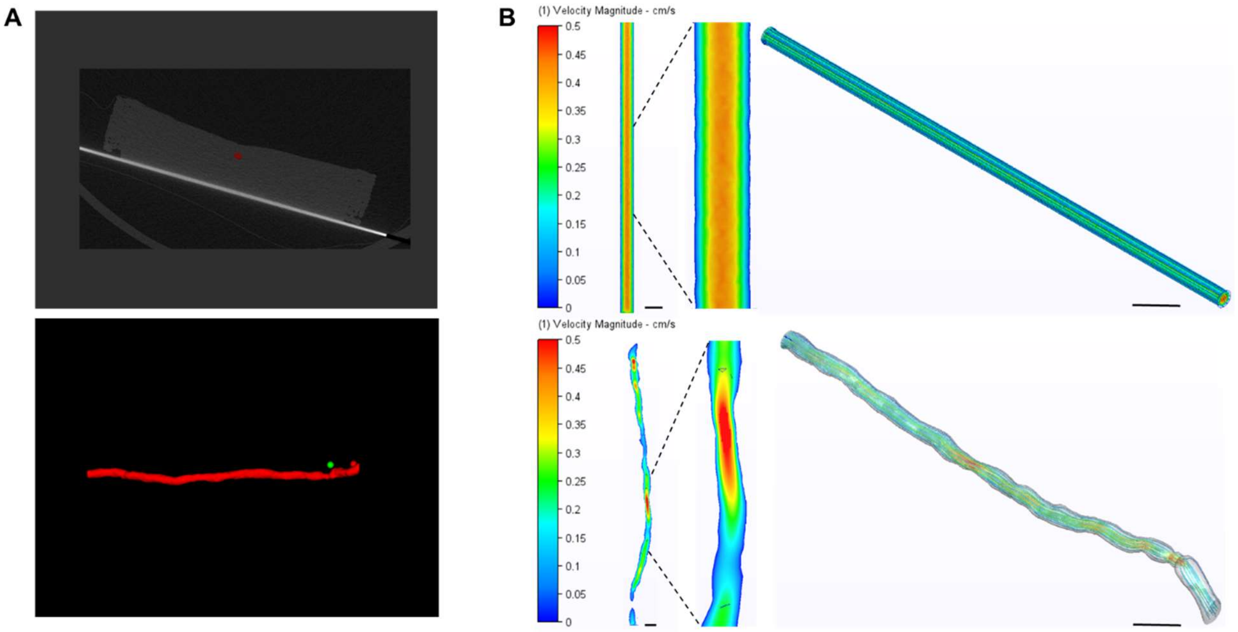


**Figure S8.** Flow analysis from straight channel CFD. a) The segmented volume of the straight channel taken from the DICOM images exported from the MicroCT. The top is the DICOM image construction with the gel and volume segmented, while the bottom shows the pulled-out volume used for analysis. b) Fluid velocity heat maps (left sides) and flow vectors (right side) for the theoretical (top) and experimental channel (bottom). Scale bars: 400μm

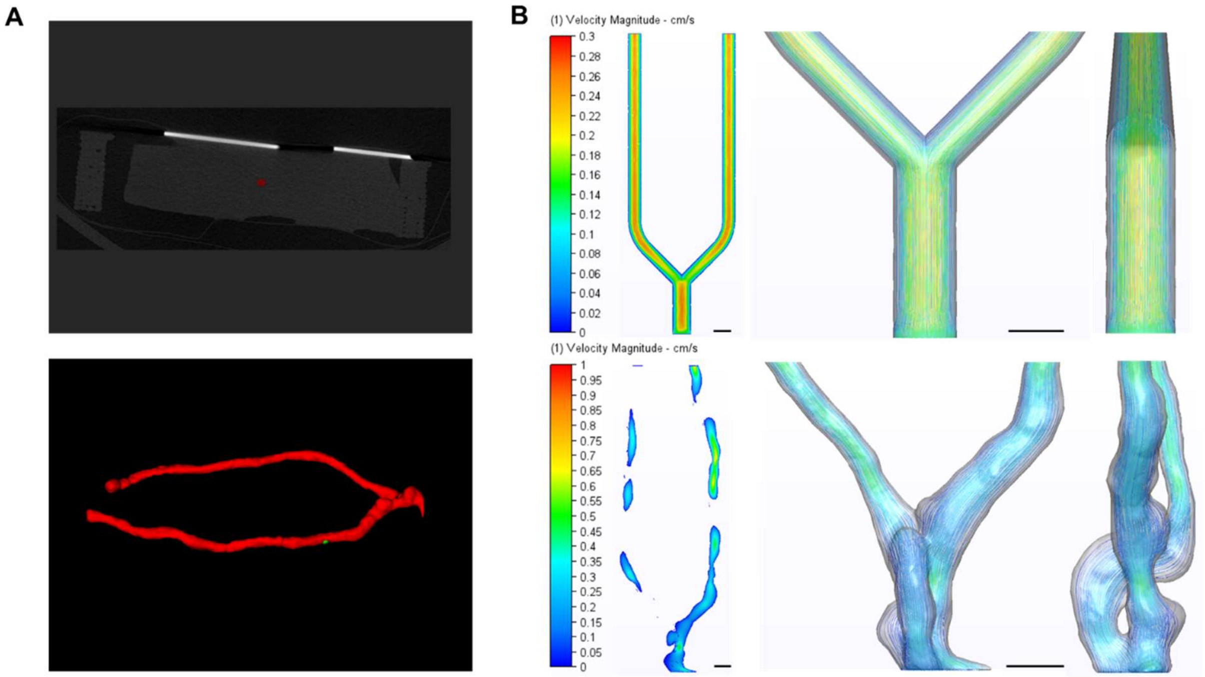


**Figure S9.** Flow analysis for Bifurcated channel CFD. a) The segmented volume of the bifurcated channel taken from the DICOM images exported from the MicroCT. The top is the DICOM image construction with the gel and volume segmented, while the bottom shows the pulled-out volume used for analysis. b) Fluid velocity heat maps (left sides) and flow vectors (right side) for the theoretical (top) and experimental channel (bottom). Scale bars: 400µm

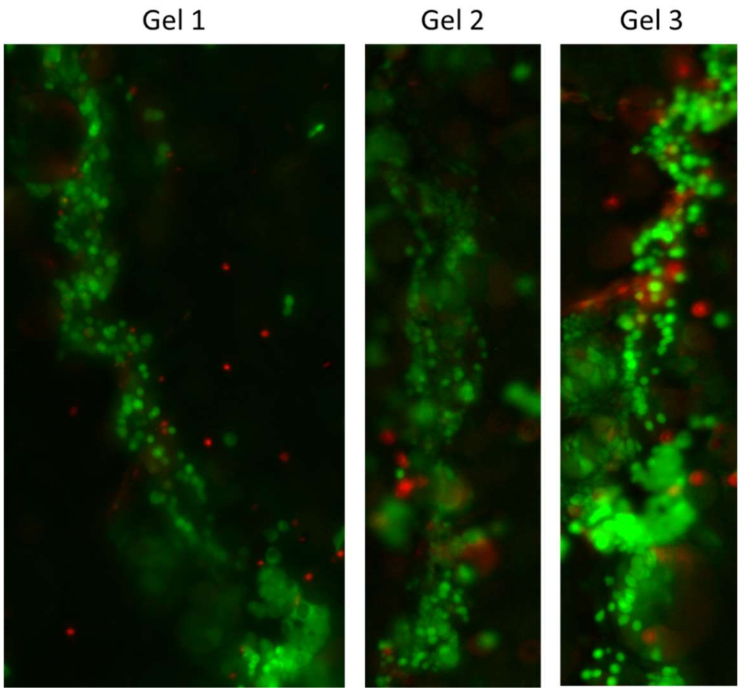


**Figure S10.** Live dead images of 3D printed B16F0 cell pellet into three suspensions with a 22G needle tip.

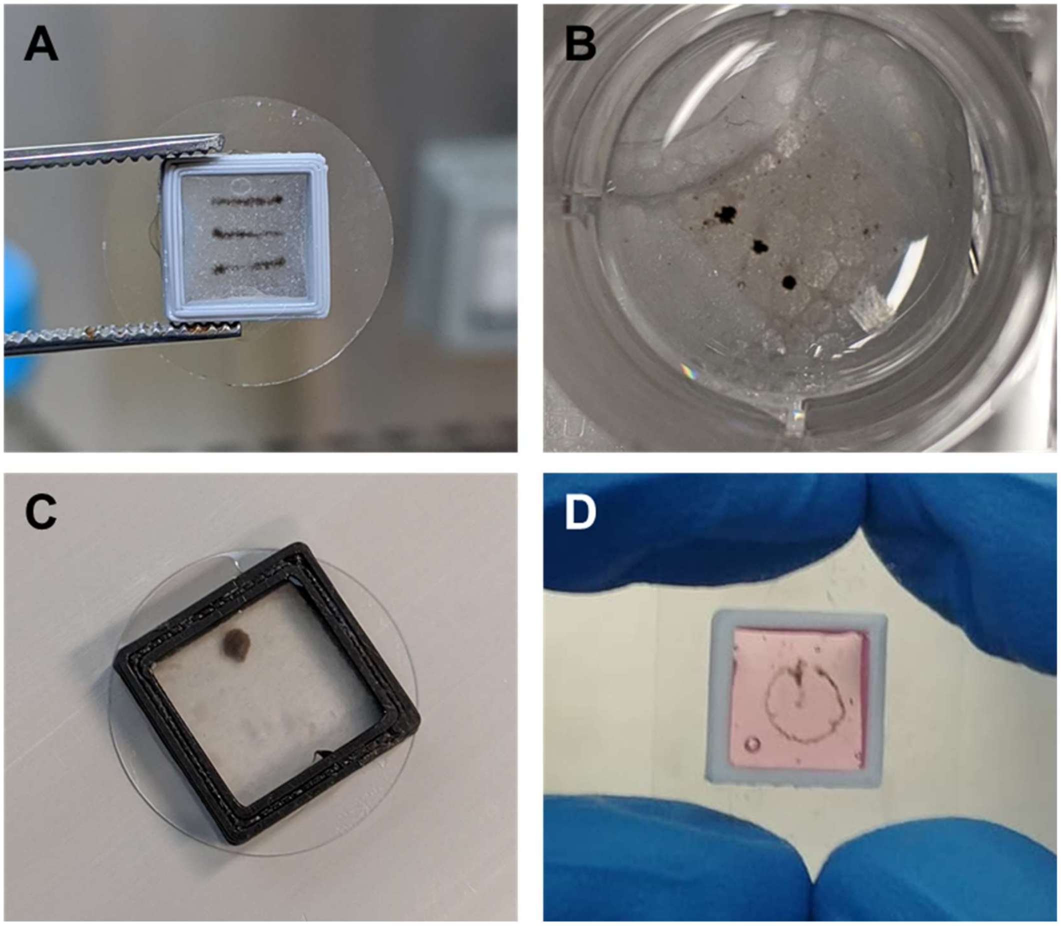


**Figure S11.** Complex Shapes of B16F0s 3D printed. Shapes shown here include three tumor lines (a), three tumor droplets (b), a 2mm diameter tumor disc (c), and a 5mm diameter tumor ring (d).

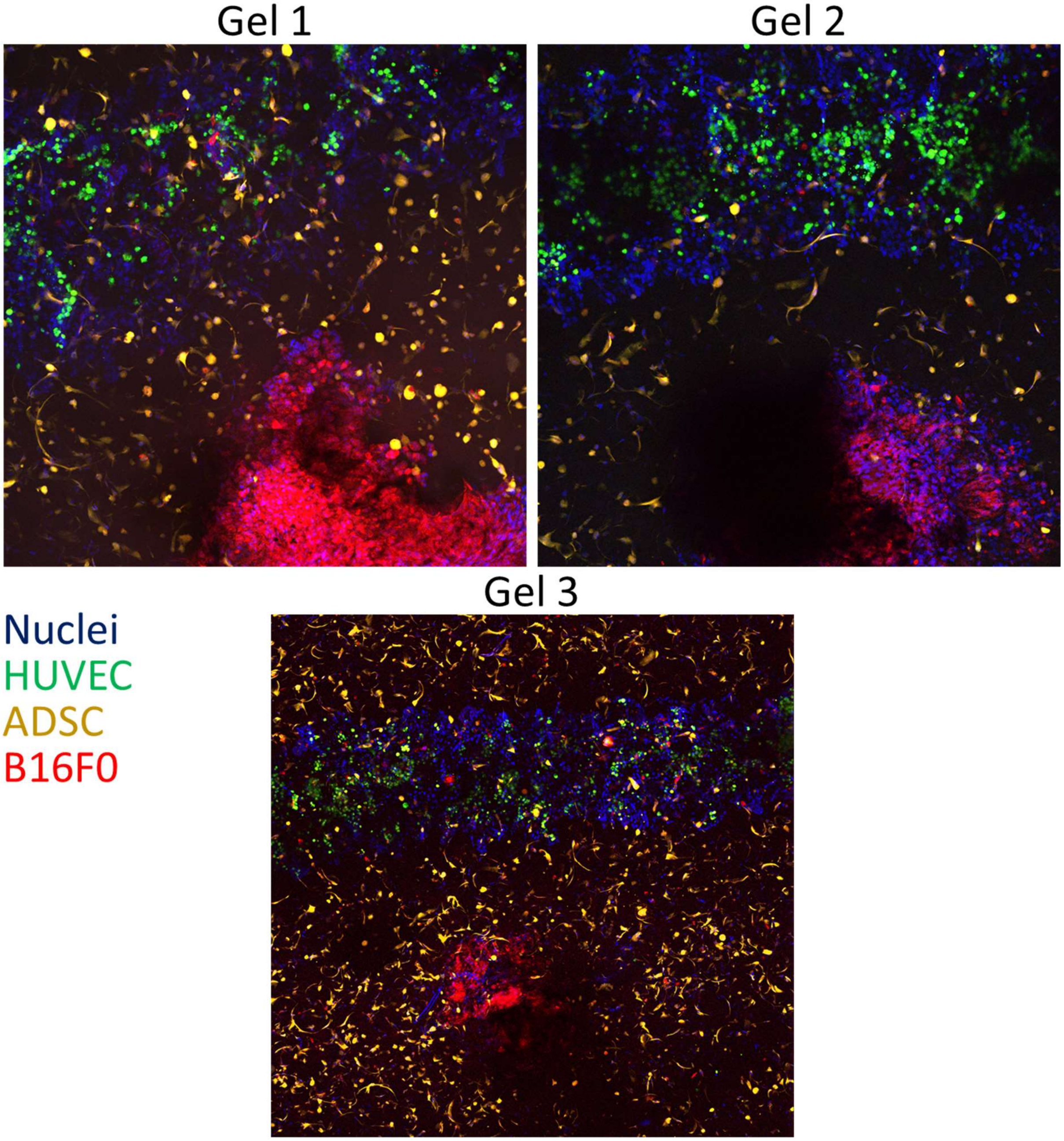


**Figure S12**. Triculture tumor model. B16F0 tumor cells (Red) migrating from printed tumor aggregate into the endothelial (green) lined channels with support ADSCs surrounding them (yellow).

NMR spectra for methacrylated gelatin samples 1 (DOF=98) and 2 (DOF=96).

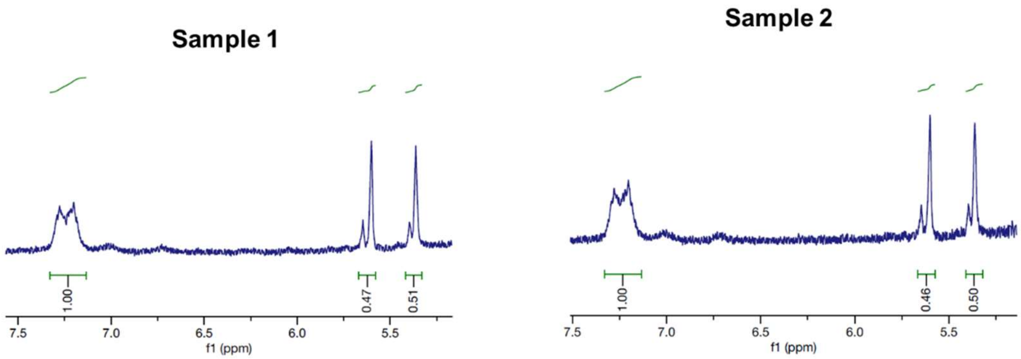


**Figure S13.** $^1$H NMR spectrum of GelMa (400.13 MHz, $D_2O$ $\delta$ 7.24 (m), 5.65 (m), 5.40 (m)) with peaks, corresponding to acrylic protons (2H) and methyl protons (3H, 1.9 ppm) from methacrylate.

| Stain | Company | Catalog # | Dilution |
| --- | --- | --- | --- |
| **Hoechst 33342** | **Life Technologies Australia Pty ltd** | **687117** | **1:200** |
| **Phalloidin-Atto 488** | **Sigma-Aldrich** | **49409** | **1:100** |
| **Rhodamine Phalloidin** | **Life Technologies Australia Pty ltd** | **R415** | **1:100** |
| **Cell tracker green** | **Life Technologies Australia Pty ltd** | **C7025** | **25µg/mL** |
| **Cell tracker red** | **Life Technologies Australia Pty ltd** | **C34552** | **25µg/mL** |
| **Cell tracker deep red** | **Life Technologies Australia Pty ltd** | **C34565** | **7.5µg/mL** |
| **Calcein** | **Life Technologies Australia Pty ltd** | **L3224** | **2 µM** |
| **Ethridium Homodimer-1** | **Life Technologies Australia Pty ltd** | **L3224** | **4 µM** |

Table S1. Information for Immunostaining
